# Cytoplasmic sequestration of the RhoA effector mDiaphanous1 by Prohibitin2 promotes muscle differentiation

**DOI:** 10.1101/283044

**Authors:** Amena Saleh, Gunasekaran Subramaniam, Swasti Raychaudhuri, Jyotsna Dhawan

## Abstract

Adhesion and growth factor dependent signalling control muscle gene expression through common effectors, coupling cytoskeletal dynamics to transcriptional activation. Earlier, we showed that mDiaphanous1, an effector of adhesion-dependent RhoA-signalling promotes MyoD expression in myoblasts, linking contractility to lineage determination. Here, we report that paradoxically, mDia1 negatively regulates MyoD function in myotubes. Knockdown of endogenous mDia1 during differentiation enhances MyoD and Myogenin expression, while over-expression of mDia1ΔN3, a RhoA-independent mutant, suppresses Myogenin promoter activity and expression. We investigated mechanisms that may counteract mDia1 to promote Myogenin expression and timely differentiation by analysing mDia1-interacting proteins. We report that mDia1 has a stage-specific interactome, including Prohibitin2, MyoD, Akt2, and β-Catenin, of which Prohibitin2 colocalises with mDia1 in cytoplasmic punctae and opposes mDia1 function in myotubes. Co-expression of mDia1-binding domains of Prohibitin2 reverses the anti-myogenic effects of mDia1ΔN3. Our results suggest that Prohibitin2 sequesters mDiaphanous1 to dampen its activity and finetune RhoA-mDiaphanous1 signalling to promote differentiation. Overall, we report that mDia1 is multi-functional signaling effector with opposing functions in different cellular stages, but is modulated by a differentiation-dependent interactome.

**Summary statement:** mDia1 has common and stage-specific functions in muscle cells. In myotubes, mDia1 is sequestered by an interacting protein Prohibitin2, which promotes Myogenin expression and mitigates mDia1’s inhibitory effects on differentiation.

**Graphical abstract:** 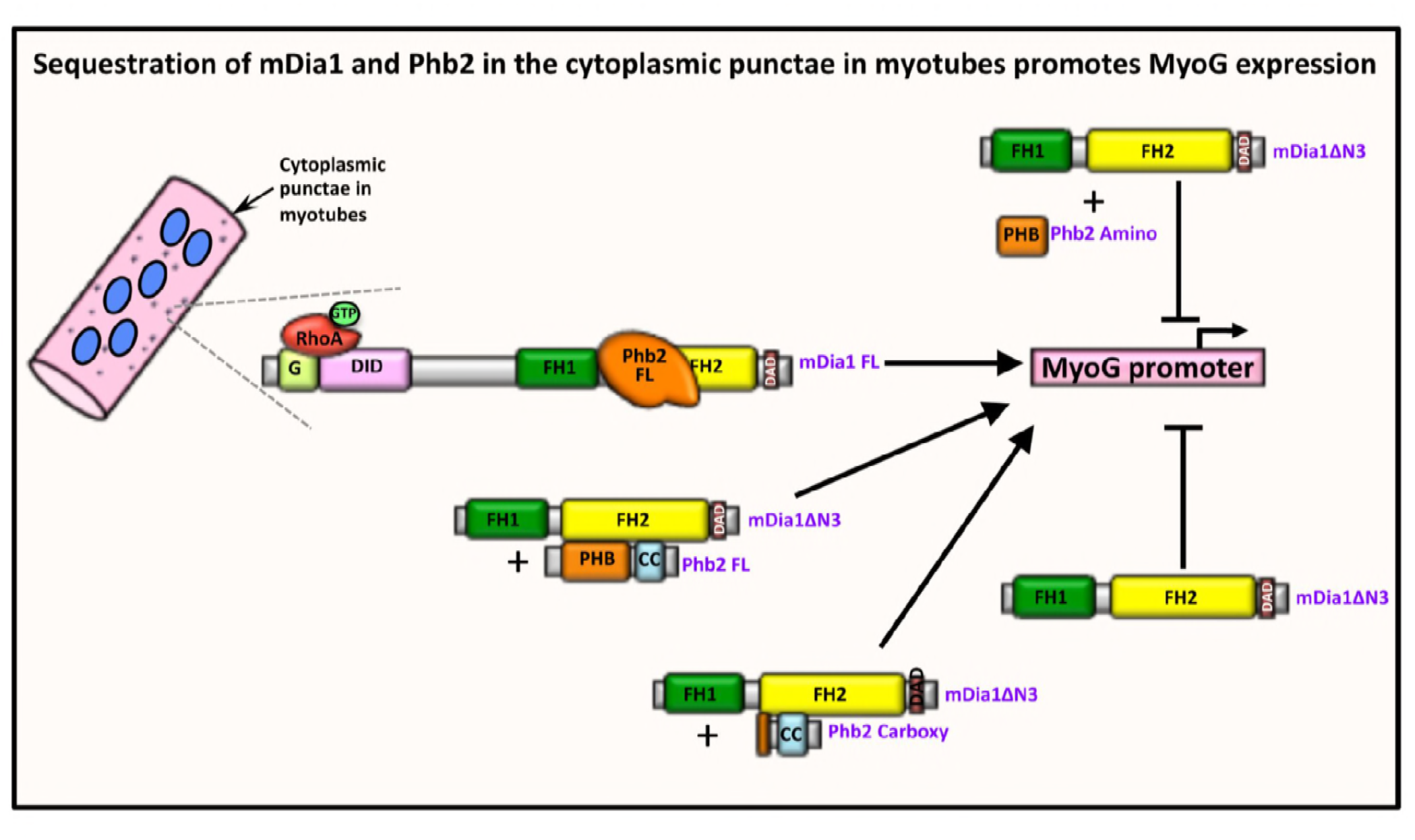

## Introduction

Cooperation between intrinsic transcriptional programs and extrinsic signaling underlies cell fate choices during development. In skeletal muscle, differentiation is regulated by muscle regulatory factors (MRFs) - MyoD, Myf5, Myogenin (MyoG) and MRF4, whose orchestrated expression and activity governs myogenic gene expression. In embryonic progenitors, MyoD and Myf5 function as lineage determinants regulating the early stages of myogenesis, whereas MyoG and MRF4 function as differentiation factors, collaborating with MEF2 to promote the later stages of myogenesis and fusion into contractile multinucleated cells (Edmondson and Olson, 1993; Rudnicki and Jaenisch, 1995; Tapscott and Weintraub, 1991). *In vitro*, myoblasts (MB) proliferate when cultured in mitogen-rich media and express MyoD, although it is transcriptionally incompetent as a consequence of growth factor-dependent post-translational modifications (Berkes and Tapscott, 2005a; Kitzmann et al., 1999; Sabourin and Rudnicki, 2001; Wei and Paterson, 2001a). Upon removal of mitogens, MyoD’s transcriptional activity is de-repressed (Lathrop et al., 1985; Massagué et al., 1986; Olson et al., 1986; Spizz et al., 1986), and MB irreversibly exit the cell cycle and fuse to form syncytial terminally differentiated myotubes (MT) (Halevy et al., 1995; Okazakit and Holtzert, 1966; Olson, 1992). In determined MB, MyoD is already engaged on muscle gene promoters genome-wide (Cao et al., 2010), but extrinsic signals are required to enhance its transcriptional competence (Tapscott, 2005; Tapscott and Weintraub, 1991), leading to activation of its key transcriptional target MyoG, and consequently, a downstream cascade of muscle-specific genes (Andres and Walsh, 1996; Faralli and Dilworth, 2012). While several signalling pathways that regulate differentiation are well known, the multiplicity of downstream effectors and mechanisms by which they channel control of muscle-specific genes is incompletely understood. In particular, the involvement of signaling mediated by cytoskeletal configuration in controlling the determination, differentiation and function of contractile muscle tissue is of interest.

Mechano-chemical cues converge with signaling by soluble growth factors such as insulin-like growth factors (IGFs) to regulate the small GTPase RhoA in myogenesis. RhoA transduces IGF and adhesion-mediated signals to control cytoskeletal dynamics that in turn impact gene expression (Welsh and Assoian, 2000). RhoA signalling is required for differentiation and its perturbation leads to reduced expression of MyoG, MRF4, MEF2 and contractile proteins (Takano et al., 1998). RhoA induces MyoD expression through regulation of actin organisation, which in turn controls the activity of Serum Response factor (SRF), a MADS box transcription factor required for MyoD expression (Carnac et al., 1998; Gauthier-Rouviere et al., 1996; L’honore, 2003; Miralles et al., 2003; Sit and Manser, 2011; Sotiropoulos et al., 1999; Soulez, 1996; Spiering and Hodgson, 2011; Wei et al., 1998). Polymerisation of globular-actin (G-actin), which sequesters Myocardin-related transcription factor (MRTF), a co-factor for SRF (Kuwahara et al., 2005), leads to the release of MRTF and its subsequent nuclear translocation to induce SRF activation (Miralles et al., 2003; Sotiropoulos et al., 1999). Ectopic expression of RhoA in proliferating MB enhances stress-fiber formation and induces the expression of differentiation-specific proteins MyoG, p21 and Troponin T (Dhawan and Helfman, 2004; Meriane et al., 2000). Interestingly, although RhoA activity is required for initial induction of myogenesis (Wei et al., 1998), its activity must be downregulated before myoblast fusion to promote myoblast fusion and differentiation (Charrasse et al., 2006; Doherty et al., 2011; Fortier et al., 2008; Nishiyama et al., 2004). Thus, actin cytoskeletal dynamics governed by RhoA signalling mediate the effects of extra-cellular stimuli to regulate MyoD expression, and play an essential role in lineage determination.

Signaling networks may have basal as well as state-specific components. The RhoA network consists of several downstream effectors (Bishop and Hall, 2000; Narumiya et al., 1997), of which mammalian Diaphanous1 (mDia1) and Rho-associated kinase (ROCK) induce actin polymerisation, and their combined actions are sufficient to mimic the effects of RhoA on focal adhesion and stress fiber formation (Amano et al., 1997; Matsui et al., 1996; Nakano et al., 1999; Wasserman, 1998; Watanabe et al., 1997; Watanabe et al., 1999). Unlike mDia1, ROCK does not mediate the effects of RhoA on MyoD expression (Dhawan and Helfman, 2004). mDia1 coordinates the dynamics of both actin filaments and microtubules (Ishizaki et al., 2001) and is known to link cytoskeletal rearrangements to transcriptional control (Copeland and Treisman, 2002; Geneste et al., 2002; Paul and Pollard, 2009; Wasserman, 1998). In proliferating MB, mDia1 functions downstream of RhoA to regulate MyoD expression by differentially modulating the activity of two transcription factors SRF and β-Catenin (Gopinath et al., 2007). However, signals emanating from the mDia1 signaling node in differentiated MT are unknown.

In this study, we probed the potential mediators of mDia1 function in myogenic cells using two screening methods to search for interacting partners. We report the interactome of this RhoA effector in MB and MT, and delineate the role of a novel myotube-specific mDia1 interacting partner Prohibitin2 (Phb2) in regulation of MyoG expression. While Dia is known to promote myoblast fusion in flies via the SCAR complex, Arp2/3 complex and actin polymerization during myofibrillogenesis (Deng et al., 2015; Deng et al., 2016), a role for mDia1 in mammalian myofibers is less well established.

The newly identified mDia1-interacting partner Phb2 (also known as Repressor of estrogen activity (REA)), is a multi-functional protein (Bavelloni et al., 2015a; Mishra et al., 2006), and is reported to regulate ERα-mediated transcription (Delage-Mourroux et al., 2000; Kurtev et al., 2004; Montano et al., 1999), CP2c-mediated transcription (Lee et al., 2008) and muscle differentiation (Héron-Milhavet et al., 2008; Sun et al., 2004). We map the domains that mediate mDia1-Phb2 interaction, identify additional signaling proteins as partners, and investigate the consequences of this interaction in regulating MyoD and MyoG expression. In summary, we report a new function for mDia1 in regulation of muscle differentiation and the protein partners that modulate this role. Our findings suggest that mDia1 plays a role in maintaining homeostatic mechanisms downstream of RhoA, with additional differentiation-dependent roles that require modulation by stage-specific interacting proteins.

## Results

### Identification of novel interacting partners of mDia1 reveals Phb2, a multi-functional transcriptional regulator

Previously we showed that mDia1 regulates the expression of MyoD, by modulating two different transcription factors-SRF and TCF-in proliferating MB (Gopinath et al., 2007). To probe the mechanisms by which mDia1 functions, we identified interacting partners for mDia1, using a yeast two-hybrid (Y2H) screen. Full-length (FL) mDia1 is auto-inhibited in the absence of active RhoA signalling (Wallar et al., 2006; Watanabe et al., 1999). To circumvent the requirement for RhoA activation in yeast we used mDia1ΔN3 (543-1192aa), a RhoA-independent constitutively active mutant of mDia1 lacking the Rho binding domain RBD (Watanabe et al., 1999) (Fig. 1A). mDia1ΔN3 fused to the GAL4 DNA-binding domain (mDia1ΔN3-BD) was used as bait, while a Matchmaker mouse cDNA library fused to the GAL4 activation domain (AD) (Clonetech), served as prey in yeast strain PJ69-4A. Putative interacting proteins for mDia1 were selected based on the induction of expression of two reporter genes – *ADE2* and *LacZ*. Prohibitin2 (Phb2) was identified as one of 8 mDia1-interacting proteins in the Y2H screen (Fig. 1B). Interestingly, Profilin1 (Pfn1), a known partner of mDia1 involved in actin nucleation (Watanabe et al., 1997) was also recovered, validating the screening strategy (Fig. S1A). Other proteins identified were all members of membrane-cytoplasmic signaling families: Niemann Pick type C2 (Npc2), Cadherin11 (Cdh11), Leukocyte receptor cluster (LRC) member8 (Leng8), Growth receptor bound protein 2 (Grb2), Protein-kinase, interferon-inducible double stranded RNA dependent inhibitor repressor of P58 (Prkrir) and Cytochrome c1 (Cyc1) (Fig. S1) (Table S1). Phb2 was selected for detailed analysis as this protein has been reported to regulate MyoD function in C2C12 MB (Héron-Milhavet et al., 2008; Sun et al., 2004). Phb2-Y2H, a flag-tagged construct encoding the partial Phb2 clone (aa 89-299) recovered in the Y2H screen (Fig. 1C), was used for co-immuno-precipitation (IP) to validate the mDia1-Phb2 interaction in mammalian cells. IP with anti-flag in HEK293T cells co-expressing GFP-tagged mDia1ΔN3 and Phb2-Y2H, resulted in co-IP of mDia1ΔN3 (Fig. 1D), confirming that ectopically expressed mDia1 and Phb2 can interact in mammalian cells.

**Figure 1.**
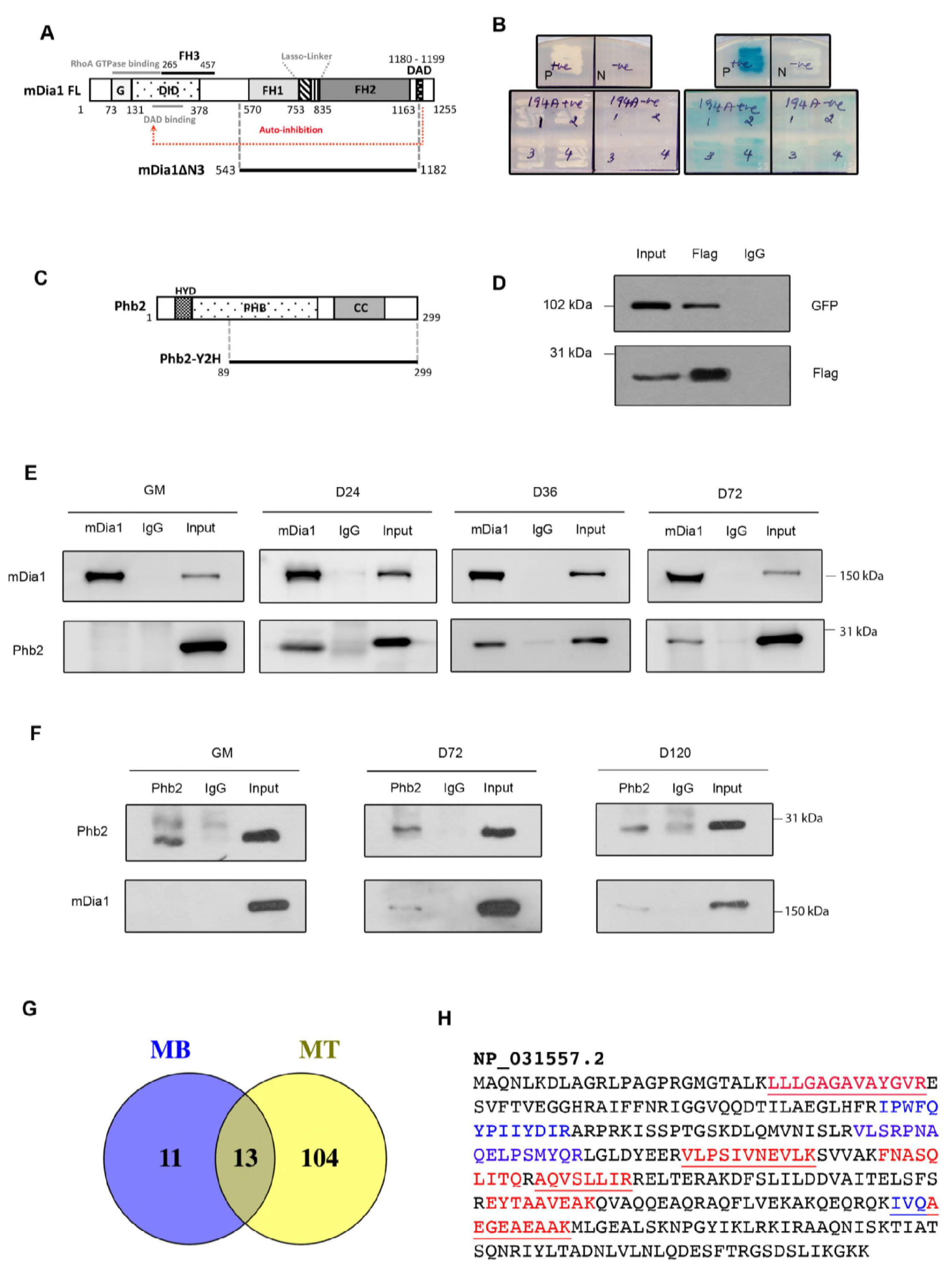
Prohibitin2, a novel mDia1-interacting protein, associates with mDia1 in myotubes. (A) Domain structure of full-length (FL) mDia1 and constitutively active mDia1 mutant, mDia1ΔN3. Grey lines indicate RhoA-GTPase and DAD binding regions. G-GTPase binding domain, DID-Diaphanous inhibitory domain, FH1, FH2, FH3-Formin Homology domains, DAD-Diaphanous Auto-inhibitory Domain. Start positions of the domains are depicted. (B) Phb2 identified as mDia1 interacting protein in a yeast two hybrid screen. PJ69-4A was co-transformed with Phb2-AD and mDia1ΔN3-BD (positive GAL4 reconstitution) or empty-BD (negative GAL4 reconstitution) and four colonies per reconstitution were screened for *ADE2* and *LacZ* reporters on -Trp/-Leu/-Ade and -Trp/-Leu+X-Gal plates respectively. Positive control “P”-*Drosophila* Batman-AD and GAGA factor-BD, negative control “N”-empty-AD and empty BD. Induction of *ADE2* is indicated by growth and induction of *LacZ* is indicated by blue colour. Trp-Tryptophan, Leu-Leucine, Ade-Adenine. AD-Activation domain, BD binding domain. (C) Domain structure of Phb2-FL and Phb2-Y2H. HYD-Hydrophobic region, PHB-Prohibitin domain, CC-Coiled coil domain. (D) Co-IP of exogenous Phb2 and mDia1ΔN3 to confirm the interaction. HEK293T, co-transfected with mDia1ΔN3 and Phb2-Y2H, and pulled down with anti-Flag antibody. (E,F) Reciprocal IP of endogenous mDia1 and Phb2 to identify stage-specific interaction. Lysates from proliferating MB (GM), MT in differentiation medium for 24 (D24), 36, (D36), 72 (D72) and 120 (D120) hours were harvested and subjected to IP with anti-mDia1 (E) or ant-Phb2 (F) antibodies. (G) LC-MS/MS analysis of mDia1-interacting proteins in myoblasts (MB) and myotubes (MT). Venn diagram represents the number of proteins that bind mDia1 in MB or MT or both MB and MT. (H) Phb2 peptides identified in MT lysates by LC-MS/MS analysis of mDia1 IP proteins. Peptides identified in the first biological replicate are indicated in red, peptides identified in the second and third replicate are shown in blue and those peptides common in all three replicates are underlined in Phb2 full-length aa sequence (NCBI Reference Sequence # NP_031557.2). The numbers represent aa position on mDia1 or Phb2 (A, C).

### Stage-specific interaction of endogenous mDia1 and Phb2 during myoblast differentiation but not in proliferation

To determine whether the interaction of mDia1 with Phb2 is relevant to muscle biology, we tested whether the endogenous proteins interact in C2C12 MB. Proliferating MB undergo differentiation when cultured in low serum, and fuse to form MT (Tapscott, 2005; Wei and Paterson, 2001b). IP was performed with whole cell lysates prepared from MB (GM-Growth medium) and MT maintained in differentiation medium for 24 (D24), 36 (D36) 72 (D72) or 120 (D120) hours (hrs). IP using anti-mDia1 antibody showed that Phb2 was specifically pulled down by mDia1 in MT (at D24-72), but not from MB (GM) (Fig. 1E). A reciprocal experiment using anti-Phb2 antibody also revealed the presence of mDia1 only in MT (upto D120) and not in MB (Fig. 1F).

To establish the expression profile of mDia1 and Phb2 during differentiation, we performed western blot analysis. The differentiation status of cultures at different time points was first established by analyzing the expression of MyoD, MyoG, Akt2 and Akt1 (Fig. S2). As expected, MyoG and Akt2 expression increased during differentiation, while MyoD and Akt1 expression decreased. Of the interacting partners, expression of Phb2 remained unchanged, whereas the expression of mDia1 decreased during differentiation. Thus, despite expression in both MB and MT, mDia1 and Phb2 interact in a stage-specific fashion only in differentiated cells.

### LC-MS/MS analysis of mDia1 interacting proteins in MB and MT

To assess the range of mDia1-interacting proteins in muscle cells, we performed LC-MS/MS analysis of mDia1-co-IPs from MB and MT. Label free quantification (LFQ) was used to identify interacting proteins and those proteins with LFQ ratio (IP/IgG) of 2 or higher were selected. mDia1 was identified in both MB and MT, confirming successful immuno-precipitation from both these states (Table S2). Notably, Phb2 was identified as an mDia1-interacting protein specifically in MT in all three replicates (Table S4), further validating the mDia1-Phb2 interaction. Phb2 peptides identified by mass spectrometry are shown in Fig. 1H. Further, 13 proteins were commonly associated with mDia1 in both MB and MT. 11 additional mDia1-interacting proteins were exclusively detected in MB and 104 were found in MT (Fig. 1G). These proteins were reproducibly detected in three independent biological replicates of mDia1 IP-LC-MS/MS analysis. Thus, mDia1 function may differ during myogenesis, with an expanded role in MT.

mDia1-interacting proteins common to MB and MT and specific to MB or MT are listed in Table S2, S3, S4 respectively. We used REVIGO (Supek et al., 2011) to perform gene ontology (GO) analysis of mDia1-interacting proteins. Proteins that associate with mDia1 in both MB and MT predominantly regulate cytoskeletal processes (Fig. S3A) and are associated with focal adhesions, cell junctions and vesicular transport (Fig. S3B). MB-specific mDia1-interacting proteins were involved in regulating cell size, nuclear import and nuclear localisation of proteins, protein folding and protein-complex assembly (Fig. S3C), and were predicted to localize to smooth endoplasmic reticulum, plasma membrane, exosomes and vesicles (Fig. S3D). However, in MT, mDia1-interacting proteins were predominantly involved in regulating multiple metabolic processes (Fig. S3E), and were predicted to localise to the cytoplasm, nucleus, mitochondria, protesome, exosomes, vesicles, extracellular compartments, ribonucleoprotein complexes, sarcomeres, focal adhesions and cell-substrate junctions (Fig. S3F).

STRING analysis of mDia1-interacting proteins was performed to identify clusters of interacting proteins in these states (Fig. S4). Associated networks of mDia1-interacting proteins common to MB and MT or specific to either MB or MT are shown (Fig. S4A-C). Interestingly, MT-specific networks of proteasomal proteins (marked red), metabolic enzymes (marked blue) and mitochondrial proteins (marked black) were identified among the mDia1-interacting proteins (Fig. S4C). Thus, the mDia1 interactome studies suggest stage-specific changes in the function of this signalling effector during myogenesis. Since Phb2 has been previously implicated in myogenic differentiation (Héron-Milhavet et al., 2008; Sun et al., 2004), we delineated the consequences of its interaction with mDia1 in detail.

### mDia1 and Phb2 co-localise in cytoplasmic punctae during differentiation

Phb2 is reported to localise to multiple cellular compartments (Mishra et al., 2006) and to function both in mitochondria and nucleus (Guan et al., 2014; Halevy et al., 1995; Massaguer; Merkwirth et al., 2008; Moncunill-massaguer et al., 2015; Montano et al., 1999; Wei et al., 2017). We evaluated the intracellular localisation of Phb2 in MB and MT by co-immunostaining Phb2 with markers of mitochondria (Cytochrome c-Cyc), cis-Golgi, (Golgi Matrix Protein of 130 kDa-GM130) or endoplasmic reticulum (Calreticulin-CALR) (Fig. S5). As in other cell types, Phb2 localized to both mitochondria and nucleus in MB and MT.

To identify the location of mDia1-Phb2 interaction we performed immuno-staining of mDia1 and Phb2 in MB and MT (Fig. 2A). Co-localisation of mDia1 and Phb2 was seen in cytoplasmic puncta in MT, but not in MB. To further evaluate the localisation of mDia1-Phb2 interaction, we used biochemical fractionation. Cytoplasmic and nuclear fractions from MT were isolated and their purity verified by western blot using antibodies against cytoplasmic markers GAPDH and nuclear markers Lamin A/C and B1 (Fig. 2B). mDia1 was predominantly cytoplasmic, with relatively lower nuclear levels whereas Phb2 was found in both nuclear as well as cytoplasmic fractions. Immuno-precipitation with anti-mDia1 antibody was performed using cytoplasmic and nuclear fractions of MT (Fig. 2C). Consistent with their co-localisation exclusively in the cytoplasm, Phb2 was co-immunoprecipitated specifically by the cytoplasmic pool of mDia1. Taken together, these findings indicate that mDia1 associates with Phb2 in cytoplasmic puncta in MT.

**Figure 2.**
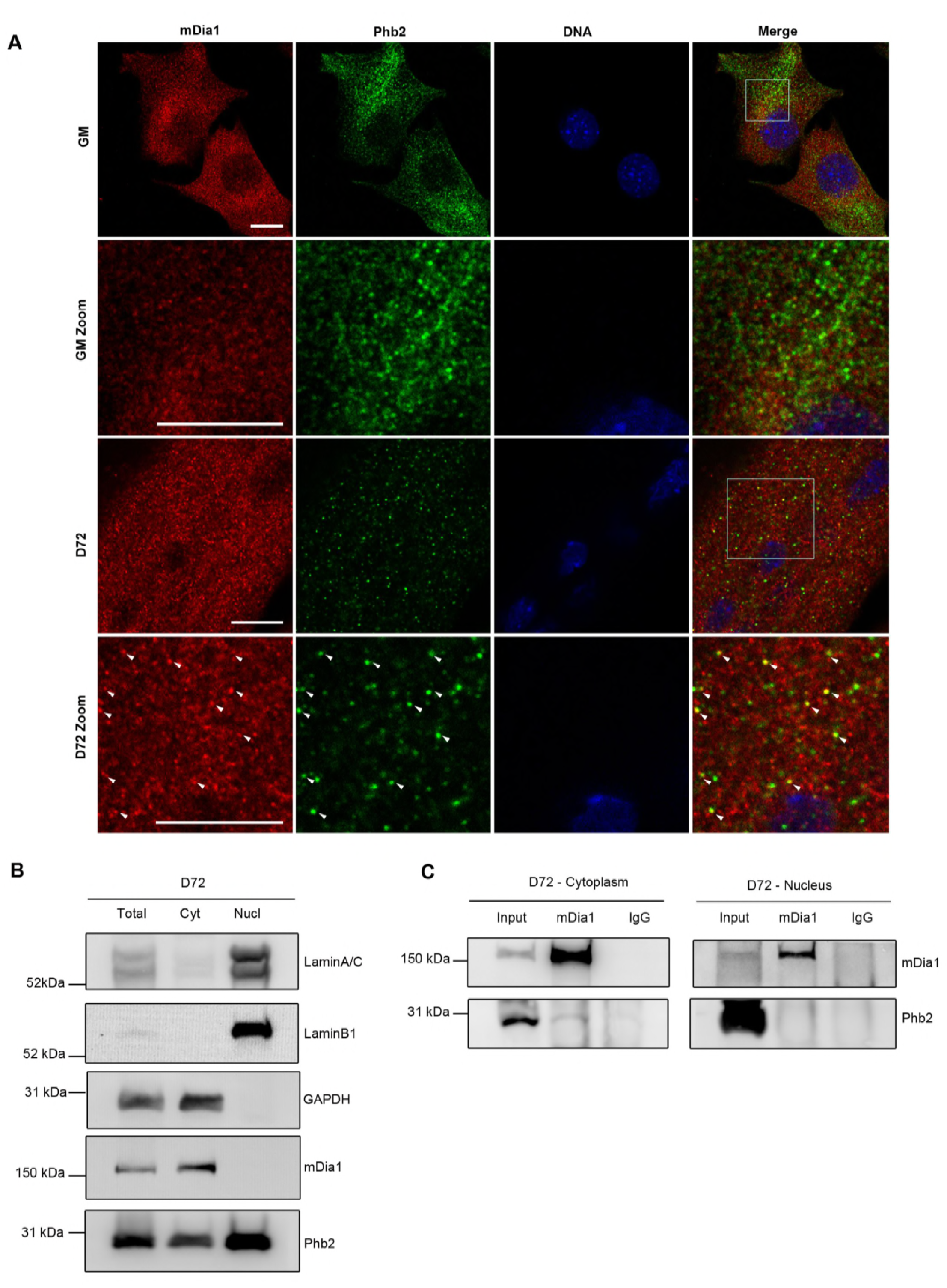
mDia1 interacts with Phb2 in the cytoplasm of myotubes. (A) Immunostaining of endogenous mDia1 and Phb2 during proliferation (GM) and differentiation (D72) to detect colocalisation. The white boxes in the merge images indicate the zoomed regions. Arrows indicate colocalised puncta. Confocal images were acquired using Leica TCS SP8 confocal microscope. Scale bar represents 10 μm. (B) Purity of cytoplasmic and nuclear fractions of MT (D72). Cytoplasmic and nuclear extracts were prepared from D72 MT, followed by analysis by western blotting with antibodies against cytoplasmic GAPDH, nuclear LaminA/C and LaminB1 to determine the purity of the fractions. Distribution of mDia1 and Phb2 was detected by western blotting using respective antibodies. (C) IP of mDia1 in Cytoplasmic and nuclear extracts to detect localisation of associated Phb2. Cytoplasmic and nuclear extracts were prepared from D72 and subjected to IP using anti-mDia1 antibody, followed by detection of Phb2. Cyt-Cytoplasm, Nucl-nucleus.

### Mapping of interaction domains on both mDia1 and Phb2

To map the interaction domains on mDia1 and Phb2 we used GFP-tagged mDia1 truncation mutants (Watanabe et al., 1999, Gopinath et al, 2007) (Fig. 3A) and flag-tagged Phb2 truncation mutants respectively (Fig. 3D). mDia1 has an N-terminal Rho Binding region including a GTPase binding region (G) and Diaphanous Inhibitory Domain (DID), three central Formin homology (FH) domains FH1, FH2 and FH3 and a C-terminal Diaphanous Autoregulatory Domain (DAD) (Maiti et al., 2012; Otomo et al., 2005; Otomo et al., 2010; Shimada et al., 2004) (Fig. 3A). In the absence of RhoA signaling, mDia1 is kept auto-inhibited through the intra-molecular interactions of its DID and DAD domains (Alberts, 2001; Lammers et al., 2005; Li and Higgs, 2003; Li and Higgs, 2005; Rose et al., 2005; Watanabe et al., 1999). Signaling from RhoA leads to release of auto-inhibition, while deletion of the aa 1-542 including the Rho binding region, results in a constitutively active mutant of mDia1, mDia1ΔN3 (Watanabe et al., 1999). The FH2 domain nucleates actin polymerisation and requires the binding of the actin binding protein Profilin1 to FH1, leading to accelerated processivity of actin filament assembly by FH1-FH2 domains (Paul and Pollard, 2009). The flexible region between FH1 and FH2 which constitutes a lasso and a linker region, is required for forming an inter-molecular ring-shaped FH2 dimer, a pre-requisite to nucleate actin polymerisation (Shimada et al., 2004; Xu et al., 2004). FH3 is a less well-defined domain which regulates the intracellular localisation of mDia1 to the mitotic spindle in HeLa cells (Kato et al., 2001).

**Figure 3.**
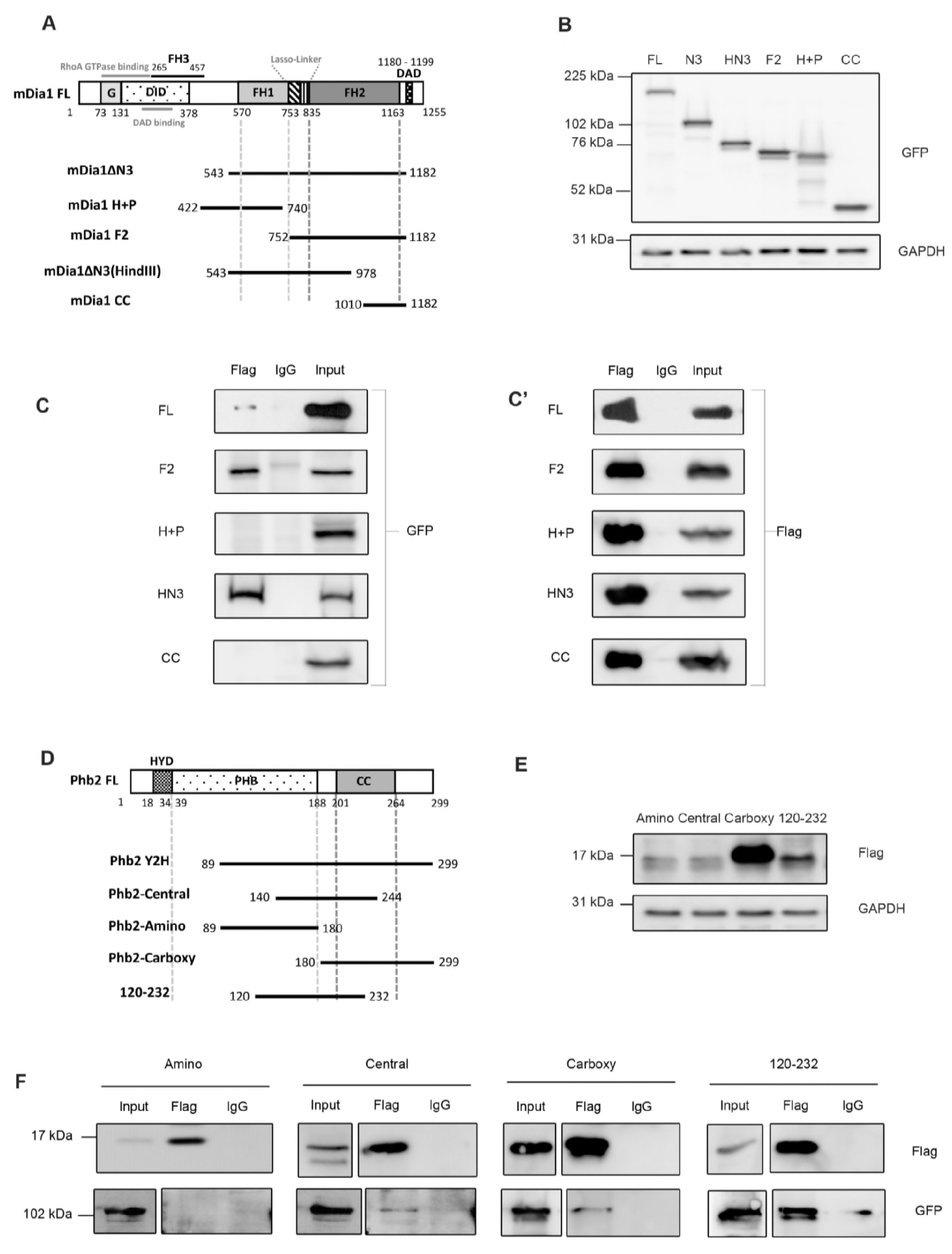
Mapping interaction domains on mDia1 and Phb2. (A) Schematic for mouse mDia1 truncation mutants. (B) Western blot to detect expression level of mDia1 mutants. Lysates from HEK293T transfected with mDia1 mutants were probed using anti-GFP antibody. GAPDH was used as a loading control. (C, C’) Co-IP of mDia1 mutants and Phb2 to map interaction domains. HEK293T cells were transfected with Phb2-Y2H and various mDia1 mutants, followed by IP with anti flag antibody. (D) Schematic for mouse Phb2 truncation mutants. (E) Western blot to detect expression of Phb2 mutants. Lysates of HEK293T transfected with Phb2 mutants were analysed using anti-flag antibody. GAPDH was used as a loading control. (F) Co-IP of Phb2 mutants and mDia1 to map interaction domains. Lysates from HEK293T cells co-transfected with various Phb2 mutants and mDia1ΔN3 were subjected to IP using anti-flag antibody. Input lanes of Phb2 Amino, Central, Carboxy and 120-232 in the GFP blot, represent lower exposures cropped from the same blot and input lanes from Central, Carboxy and 120-232 in flag blot represent higher exposures cropped from the same blot. The numbers represent aa positions (A, D).

Phb2 contains an N-terminal hydrophobic single trans-membrane alpha helix (aa 18-34) (HYD), a central Prohibitin (PHB) domain (aa 39-201) and a C terminal Coiled coil (CC) domain (aa 188-264) (Merkwirth and Langer, 2009) (Fig.2D). Human and mouse Phb2 proteins are 100% identical. Other important sequences include a positively charged N terminal leader sequence (aa 1-50) that functions as a mitochondrial targeting sequence (MTS) in both human and mouse Phb2 and a nuclear localization signal at the C terminal in the human Phb2 (Kasashima et al., 2006; Merkwirth et al., 2008). Putative nuclear localisation signal (aa 86-89) and nuclear receptor box have been predicted for mouse Phb2 (Merkwirth et al., 2008). The central PHB domain is predicted to facilitate oligomerisation of Phb1/2 and may also facilitate membrane association and partitioning into lipid micro-domains (Mishra et al., 2006; Morrow and Parton, 2005; Winter et al., 2007), whereas the coiled coil domain (aa 190-264) promotes the formation of large ring-like oligomeric complexes of Phb1 and Phb2 in the mitochondrial membrane (Merkwirth and Langer, 2009; Tatsuta et al., 2005). Flag-tagged truncation mutants of Phb2 were generated to span aa 89-299 of Phb2-FL (full-length), the region recovered in the Y2H screen (Fig. 2D).

To map the regions of mDia1 that interact with Phb2, HEK293T cells were co-transfected with flag-tagged Phb2-Y2H (aa 89-299) and different mDia1 truncation mutants (GFP-tagged). Western blotting of mDia1 and Phb2 truncation mutants revealed that the mDia1 mutants expressed at relatively equal levels whereas the Phb2-Carboxy mutant expressed at a higher level than the other weakly-expressing Phb2 mutants (Fig. 3B, Fig. 3E respectively). Co-IP of HEK293T lysates using anti-flag antibody was followed by detection with anti-GFP antibody (Fig. 3C,C’). mDia1ΔN3, mDia1F2, mDia1ΔN3(HindIII) interacted with Phb2-Y2H, whereas mDia1H+P and mDia1CC did not. This analysis indicates that mDia1 binds Phb2-Y2H via aa 752-978, which maps to the lasso-linker region between the FH1-FH2 domains including a portion of the FH2 domain. We previously reported that the FH1 domain mediates SRF-independent regulation of MyoD whereas the FH2 domain mediates SRF-dependent regulation of MyoD expression (Gopinath et al., 2007). Conceivably, binding of Phb2 to this region (aa 752-978) between FH1 domain and FH2 domain might contribute to MyoD regulation.

Reciprocally, to map the regions of Phb2 that interact with mDia1, HEK293T cells were co-transfected with different flag-tagged Phb2 truncation constructs and mDia1ΔN3-GFP, followed by IP with anti-flag antibody (Fig. 3F). Phb2 Central (aa 140-244), Carboxy (aa 180-299), and 120-232 (aa 120-232) region bound mDia1, however the Amino (aa 89-180) region did not. This analysis revealed that the minimal mDia1-interacting region of Phb2 maps to aa 180-232. This region includes a small portion of the PHB domain and almost half of the coiled coil domain and lies within aa 120-232 region of Phb2. The region aa 120-232 of human PHB2 contains overlapping binding sites for MyoD, Akt2 and Estrogen receptor α (ERα) (aa175-198) (Delage-Mourroux et al., 2000; Sun et al., 2004) (Table S5). Since mouse and human Phb2 proteins are 100% identical, we infer that the binding of mouse mDia1 to Phb2 could compete with the binding of Phb2 to Akt2, MyoD or ERα. Taken together, these domain-mapping studies suggest the possibility that Phb2 may regulate gene expression by forming mutually exclusive interactions with key transcription regulators and effector proteins.

### Phb2 forms a complex with mDia1 and pro-myogenic proteins during differentiation

To determine whether mDia1 and Phb2 formed additional interactions with known muscle transcriptional regulators, we pulled down mDia1 and probed for co-immunoprecipitation of Akt2, MyoD and β-Catenin. mDia1 also associated with Akt2 and pAkt2(ser474) specifically in MT (Fig. 4A). MyoD was also found to interact with mDia1 only during differentiation, along with Phb2 and Akt2 (Fig. 4B). Further, active β-Catenin was pulled down with mDia1 along with Phb2 (Fig. 4C). Interestingly, mDia1 also co-immuno-precipitated the transcriptional regulator Phb1, a known partner of Phb2 (Kasashima et al., 2006; Mishra et al., 2005), suggesting a role for this complex in gene regulation (Fig. 4D). Akt2 and Phb2 were also co-immunoprecipitated by mDia1 along with Phb1. These findings indicate the existence of multi-protein complexes that contain mDia1 and differentiation-regulating proteins specifically in MT. Taken together, our findings suggest that the mDia1-Phb2 protein complex might associate with one or more of the mDia1-interacting partners pAkt2, MyoD and active β-Catenin to regulate differentiation.

**Figure 4.**
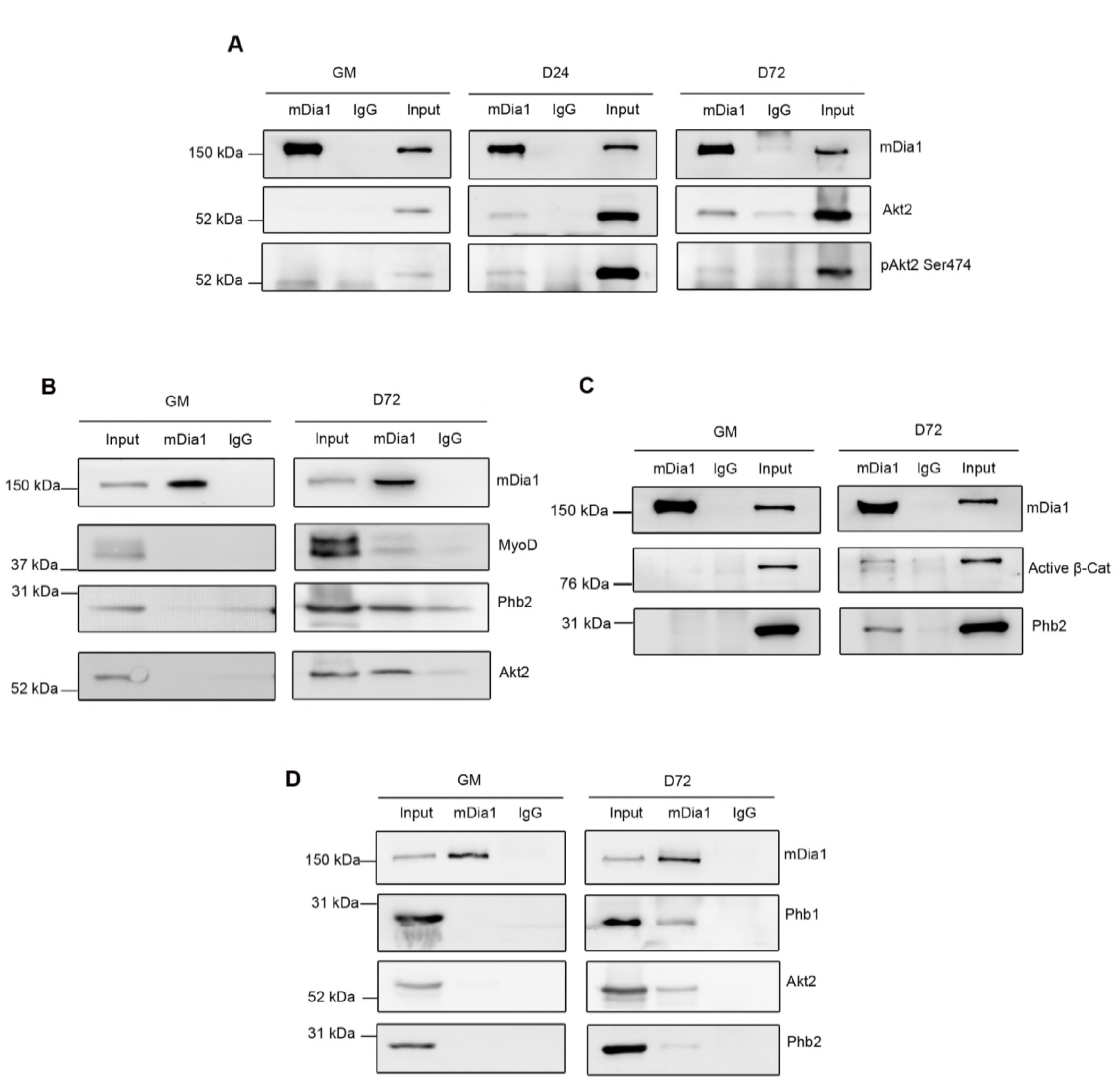
mDia1 co-immunoprecipitates differentiation markers MyoD, active β-Catenin and pAkt2 (Ser474) along with Phb2 during differentiation. C2C12 cells cultured under growth conditions (GM) or differentiated for 24 (D24) or 72 (D72) hours were lysed and subjected to IP with anti-mDia1 antibody, and analysed by western blotting using respective antibodies. (A) Co-IP of mDia1 with Akt2 and pAkt2(ser474) in MT. (B) Co-IP of MyoD, Phb2 and Akt2 by mDia1 in MT. (C) Co-IP of active β-Catenin and Phb2 by mDia1 in MT. (D) IP of Phb1, Akt2 and Phb2 by mDia1 in MT. Act β-Cat-Active β-Catenin.

### Endogenous mDia1 represses MyoD and MyoG expression

To determine the functional significance of the mDia1-Phb2 interaction in MT, mDia1 and Phb2 were knocked down using siRNA SMART pools (each comprising 4 independent siRNAs). Briefly, MB were transfected with scrambled control (SCR), mDia1, Phb2 or mDia1+Phb2 siRNA pools and shifted to DM for 48h. RNA was isolated from the knockdown samples and transcript levels of mDia1, Phb2, MyoD and MyoG were evaluated using quantitative reverse transcription PCR (qRT-PCR) (Fig. 5A). mDia1 transcripts were reduced to 31% and 40% in mDia1 and mDia1+Phb2 knockdown MT respectively. Phb2 transcripts were reduced to 45% and 53% in Phb2 and mDia1+Phb2 knockdown MT respectively. There was a small but significant increase in MyoD and MyoG transcript levels when mDia1 or mDia1+Phb2 were knocked down, however knockdown of Phb2 alone did not affect MyoD or MyoG transcript expression. These observations suggest that endogenous mDia1 mildly represses MyoD and MyoG expression in MT, whereas Phb2 on its own does not affect their expression.

**Figure 5.**
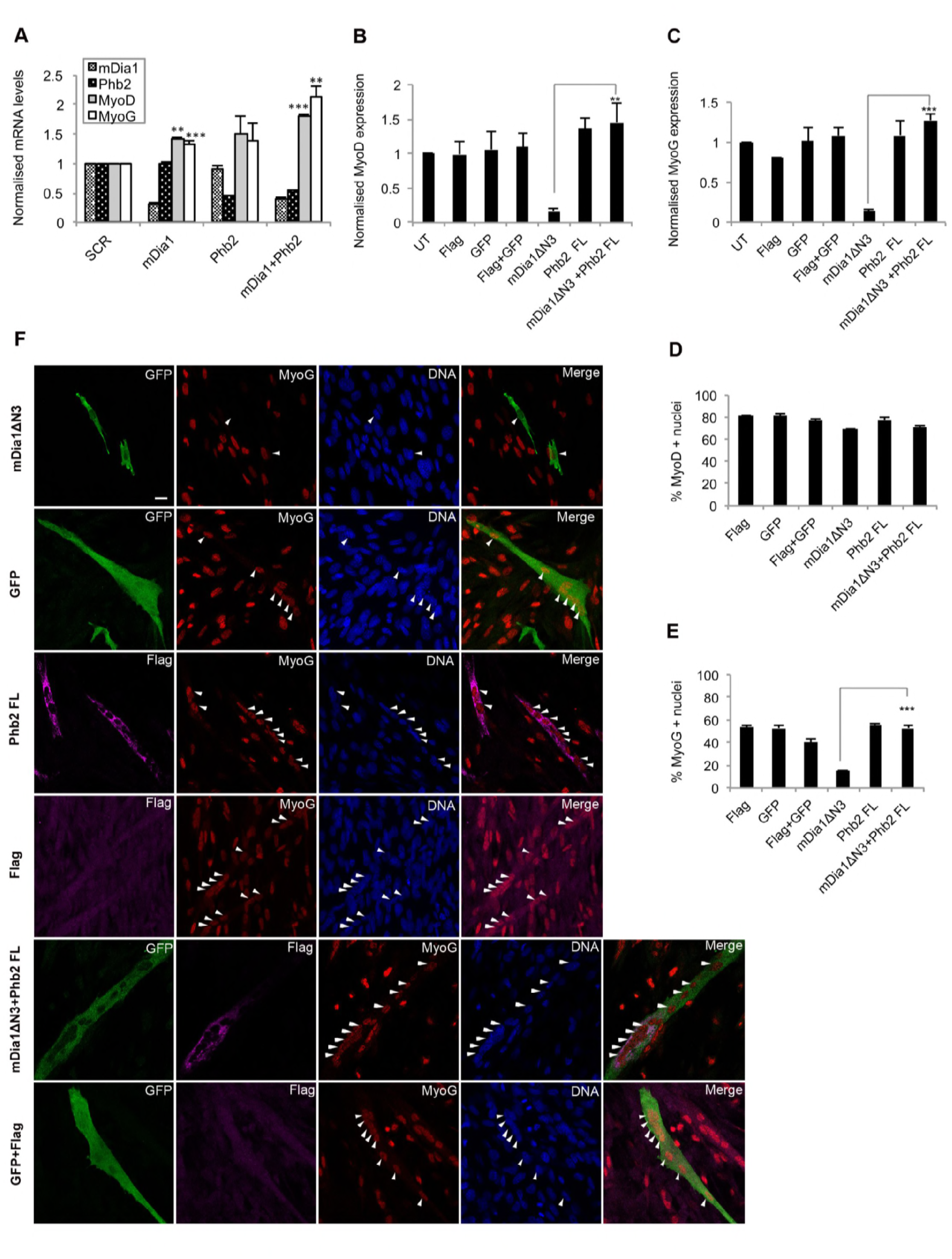
Co-expression of mDia1 and Phb2 prevents repression of endogenous MyoG and MyoD. (A) Knockdown of mDia1 (but not Phb2) up-regulates MyoD and MyoG. qRT-PCR analysis of MyoD and MyoG transcripts in mDia1 and Phb2 knockdown in MT. C2C12 MB were transfected with control scrambled (SCR), mDia1, Phb2 or both mDia1+Phb2 siRNA pools and shifted to DM for 48 hours, followed by RNA extraction and qRT-PCR analysis. **p<0.01, ***p <0.001 compared with control SCR, n=3. Bar graph represents respective mRNA values normalized to GAPDH and SCR control. Error bars represent ± s.e.m. (B, C) Overexpressed mDia1ΔN3 represses MyoD and MyoG in MT, while co-expressed Phb2 reverses the repression. qRT-PCR analysis of MyoD (B) and MyoG (C) transcripts respectively in C2C12 transiently transfected with mDia1ΔN3 and Phb2 and shifted to DM for 36 hours. ***p<0.001, p<0.01, n=3. Bar graphs indicate normalised mRNA values. UT and GAPDH were used for normalisation of mRNA levels. Error bars represent ± s.e.m. (D, E) MyoG protein (but not MyoD) is repressed by mDia1ΔN3. Immunostaining of endogenous MyoD and MyoG in MT ectopically expressing mDia1 and Phb2. C2C12 were transfected with mDia1ΔN3 and Phb2-FL and shifted to DM for 36 hours, followed by IFA for Flag (Phb2), GFP (mDia1), MyoD and MyoG. Percentage of MyoD (D) and MyoG (E) positive nuclei were determined by counting atleast 200 cells. ***p<0.0001, n=3. (F) Representative images of endogenous MyoG protein during over-expression of mDia1ΔN3 and Phb2 during differentiation. Scale bar represents 20 μm. UT-Untransfected, FL-Full-length.

### Over-expression of mDia1 leads to repression of Myogenin, which is reversed by co-expressed Phb2

To further analyze the role of mDia1 and Phb2 in regulating MyoD and MyoG expression, we performed over-expression studies in MT (Fig. 5B,C respectively). While expression of exogenous Phb2-flag did not affect either transcript, expression of mDia1ΔN3 strongly suppressed the level of MyoD and MyoG transcripts to 17% and 13% of control respectively. Notably, when Phb2 was co-expressed with mDia1ΔN3, MyoD and MyoG mRNA levels were restored to control levels, suggesting that this interacting protein counteracts mDia1’s repressive function. To further assess the effect of ectopic Phb2 and mDia1ΔN3 on MyoD and MyoG protein expression in MT, we used immunofluorescence. Ectopic expression of mDia1ΔN3 and Phb2-FL did not affect the number of MyoD+ cells (Fig. 5D). mDia1ΔN3 expression alone reduced the number of MyoG+ cells to 15%, whereas 55% of Phb2-FL expressing cells were MyoG+. Interestingly, co-expression of Phb2 with mDia1ΔN3 restored the frequency of MyoG+ cells to 53%, comparable to control (Fig. 5E,F). Taken together, these findings suggest that exogenous mDia1 represses MyoD and MyoG mRNA expression at the level of mRNA, but only MyoG protein levels were affected. Further, Phb2 may function in MT to block mDia1’s repressive effect on MyoG.

### Co-expression of mDia1ΔN3 and Phb2 relieves the repression of MyoG promoter

To evaluate the functional significance of mDia1 and Phb2 interaction, we studied the effect of ectopically expressed mDia1ΔN3 and Phb2 on MyoG transcription. A 1565 bp region of the MyoG promoter containing E-boxes and other regulatory elements controls MyoG expression (Edmondson et al., 1992a). We used promoter-reporter assays where MyoG promoter-luc constructs were transfected into C2C12 cells along with individual mDia1 mutants and Phb2-FL during differentiation. The MyoD promoter (DRR-luc) was not regulated by the mDia1-Phb2 interaction (data not shown). However, in MyoG promoter assays in MT, mDia1ΔN3 individually reduced the activity of the MyoG promoter whereas mDia1H+P and Phb2 did not (Fig. 6A). Interestingly, as with endogenous MyoG expression, when Phb2 was co-expressed with mDia1ΔN3, MyoG promoter activity returned to control levels. Co-expression of Phb2 with mDia1H+P or mDia1CC mutants that do not interact with Phb2, did not affect MyoG promoter activity. Thus, mDia1ΔN3-Phb2 interaction is required to rescue mDia1-mediated repression of MyoG promoter activity.

**Figure 6.**
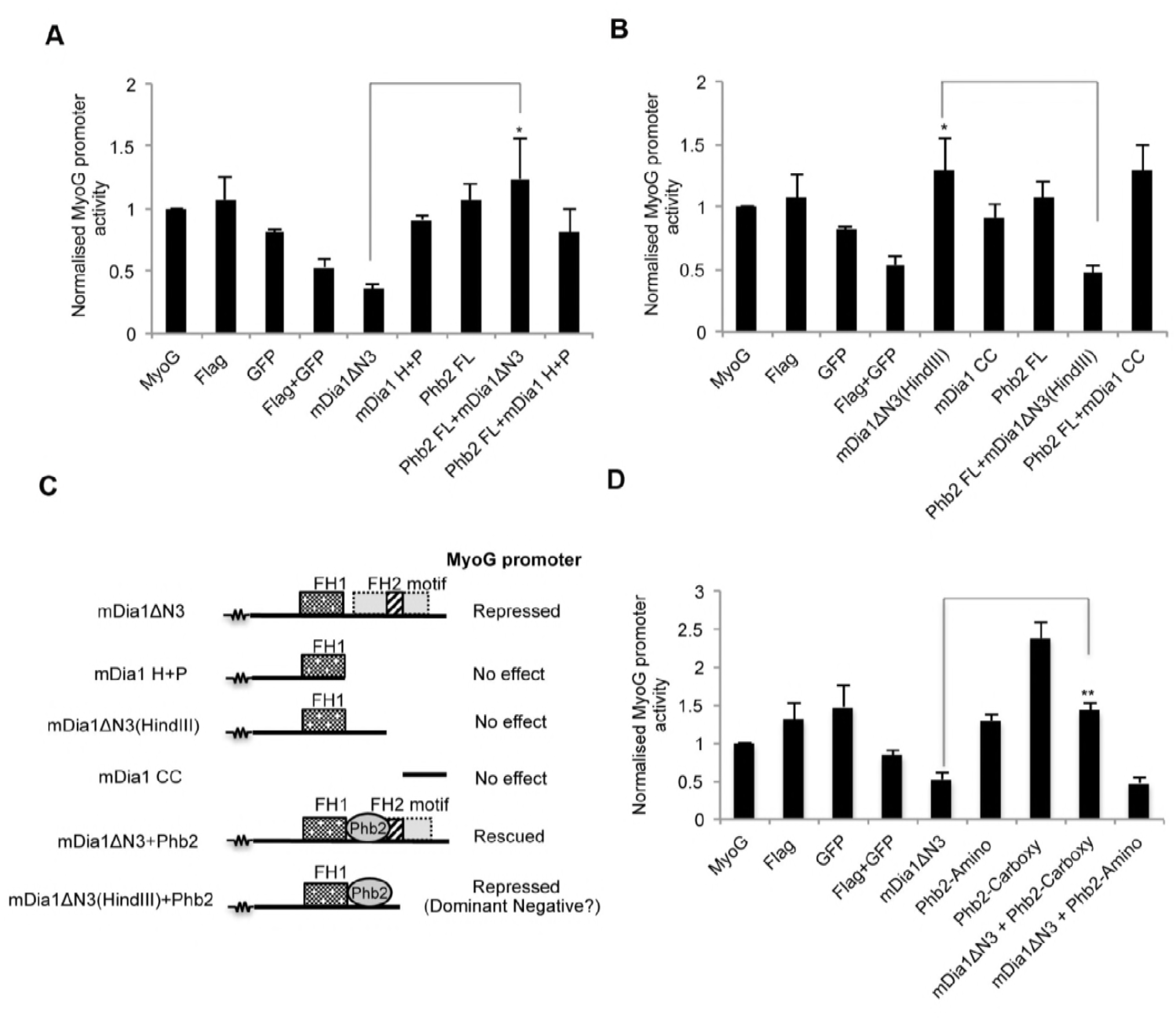
Co-expression of mDia1ΔN3 and Phb2 rescues MyoG promoter activity. C2C12 were transfected with various mDia1 and Phb2 mutants along with MyoG-promoter reporter construct and shifted to DM for 72 hrs, followed by lysis and dual-luciferase assays. (A) Normalised MyoG promoter activity in MT transfected with mDia1ΔN3, mDia1H+P or Phb2-FL. *p<0.05, n=3. (B) Normalised MyoG promoter activity in MT transfected with mDia1ΔN3(HindIII), mDia1CC or Phb2-FL. *p<0.05, n=3. (C) Schematic illustrating FH2 motif (aa 946-1010)-mediated regulation of MyoG promoter by mDia1 mutants and Phb2. The squiggle represents the common domains not depicted. The FH2 motif is indicated by the stripped box within the dotted grey box representing the FH2 domain. (D) Normalised MyoG promoter activity in MT transfected with mDia1ΔN3, Phb2 carboxy or Phb2 amino. **p<0.01, n=3. For all Luciferase assays performed, Luciferase readings were normalised to Renilla Luciferase, empty pGL3 vector and basal DRR or MyoG promoter activity, to correct for background luminescence and transfection efficiency. Bar graphs represent normalised Luciferase values. Error bars represent ± s.e.m. FL-Full-length. UT-Untransfected.

Another Phb2-interacting mDia1 mutant, mDia1ΔN3(HindIII) when expressed alone did not affect MyoG promoter activity (Fig. 6B), as with Phb2 alone. Unexpectedly, co-expression of the interacting pair Phb2 and mDia1ΔN3(HindIII) reduced the MyoG promoter activity. This data suggests that the mDia1ΔN3(HindIII)-Phb2 interaction represses MyoG activity. mDia1 mutants that include the FH2 subdomain-FH2 motif (aa 946-1010) that lies within the FH2 domain (Shimada et al., 2004), repressed MyoG promoter activity (Fig. 6C, Table 1). Notably, Phb2’s interaction with mDia1ΔN3, which includes FH2-motif, rescued the repression of MyoG promoter while its interaction with mDia1ΔN3(HindIII), which lacks FH2-motif repressed the MyoG promoter. Taken together, these findings indicate that mDia1ΔN3-Phb2 interaction is required to rescue MyoG promoter activity and there are additional FH2-motif specific mechanisms operating to regulate MyoG promoter.

**Table 1.** Regulation of MyoG promoter by mDia1 mutant-Phb2 interaction.

| Expressed proteins | FH2 motif | MyoG promoter | Proposed mechanism of action |
| --- | --- | --- | --- |
| mDia1ΔN3 | Present | Repressed | FH2 motif-mediated repression |
| mDia1H+P | Absent | No effect | No FH2 motif-mediated repression |
| mDia1ΔN3(HindIII) | Absent | No effect | No FH2 motif-mediated repression |
| mDia1CC | Absent | No effect | No FH2 motif-mediated repression |
| Phb2 | Not applicable | No effect | No effect |
| mDia1ΔN3+Phb2 | Present in mDia1 mutant | Rescued | Phb2 recruits pro-myogenic proteins to FH2 motif |
| mDia1ΔN3(HindIII)+Phb2 | Absent in mDia1 mutant | Repressed | Acts as dominant negative (Phb2 cannot recruit pro-myogenic proteins to FH2 motif, since mDia1ΔN3(HindIII) lacks FH2 motif) |

To further delineate Phb2’s rescue of MyoG promoter activity from repression by mDia1ΔN3, we transfected mDia1ΔN3 along with different Phb2 truncation constructs. As seen earlier, mDia1ΔN3 on its own repressed MyoG promoter activity (Fig. 6D). Consistent with the results using different mDia1 domains, MyoG promoter activity was rescued by co-expression of mDia1-interacting Phb2 mutant, Phb2-Carboxy, but not by co-expressing Phb2-Amino, a mutant that does not interact with mDia1. These findings indicate that only the Phb2-Carboxy mutant that included the mDia1-interacting region rescued the MyoG promoter activity, while Phb2-Amino that lacked the mDia1-interacting region did not. On its own, the interacting Phb2-Carboxy mutant induced MyoG promoter activity, consistent with sequestering endogenous mDia1, while the non-interacting Phb2-Amino did not. Together, these findings emphasize a role for Phb2 in mitigating the repressive effect of mDia1 on MyoG promoter activity in MT.

In summary, we report that mDia1 is involved in differentiation-specific interactions with multiple transcriptional regulators Phb2, MyoD, pAkt2 and active β-Catenin, suggesting the involvement of one or more complexes of signalling molecules and transcription factors focused on control of MyoG expression (Fig. 7). Both endogenous and exogenously expressed mDia1 repress MyoG expression at the transcript as well as protein level. However, when bound to Phb2, mDia1 does not repress MyoG, suggesting that the differentiation-specific interaction of mDia1-Phb2 is required to block mDia1-mediated repression of MyoG. Moreover, the mDia1-Phb2 interaction was localized to cytoplasmic puncta in MT, indicating that Phb2 may sequester mDia1 to regulate its activity and mitigate repression of MyoG. Taken together with our earlier report of RhoA-mDia1 signaling impact on MyoD expression in undifferentiated MB, these findings suggest that adhesion/contractility-dependent signaling circuits also control differentiation.

**Figure 7.**
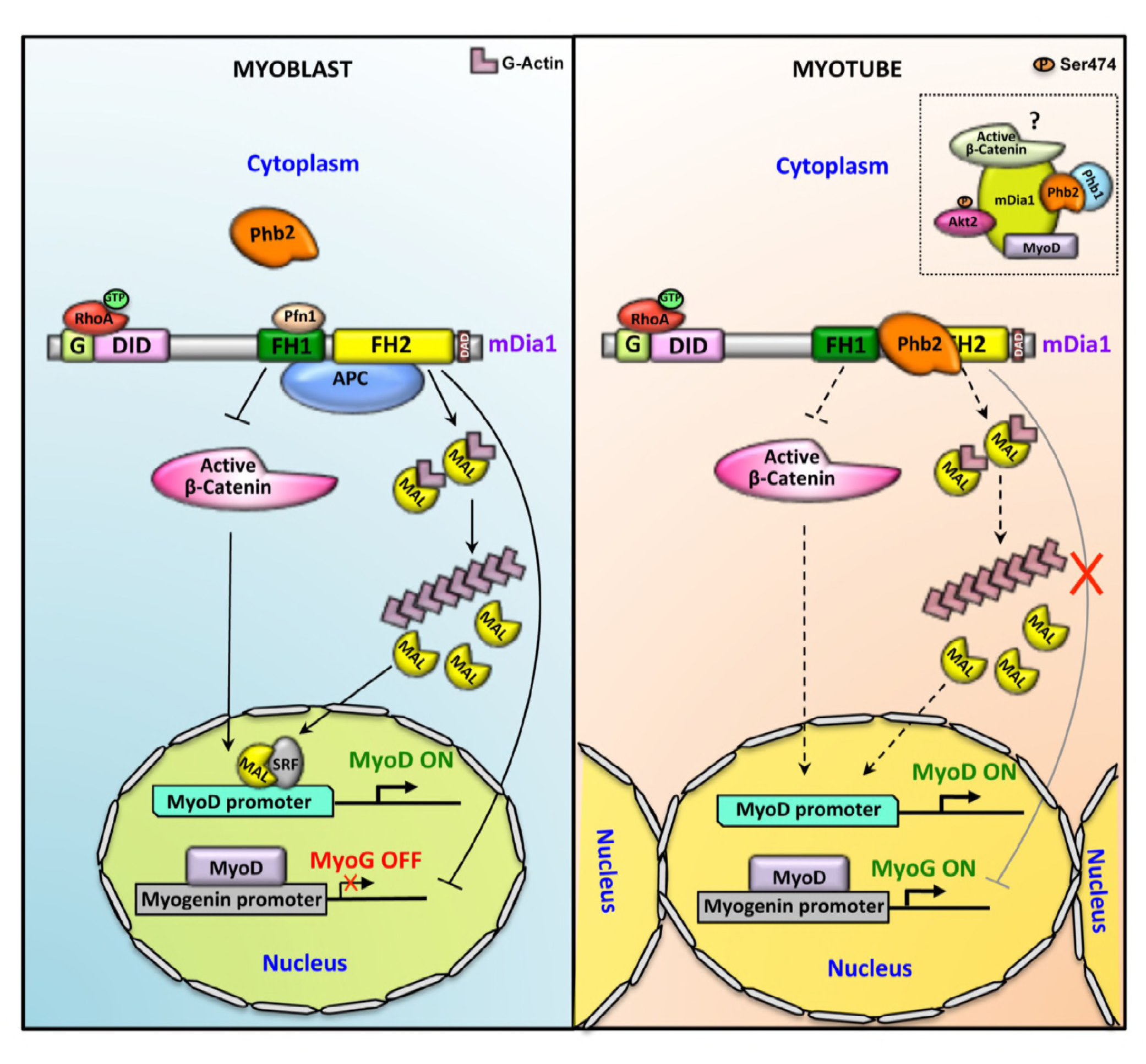
Model: Phb2 sequesters mDia1 in the cytoplasmic puncta during differentiation to promote MyoD function. In MB, mDia1 regulates MyoD expression by titrating the activity of two antagonistic pathways involving β-Catenin and SRF. Binding of APC to the region between FH1-FH2 (Wen et al., 2004) domains prevents nuclear localisation of β-Catenin and inhibits MyoD expression (Gopinath et al., 2007). In the absence of Phb2 binding, the FH2 motif within the FH2 domain of mDia1 inhibits MyoG expression. In MT, Phb2 binds and sequesters mDia1 in the cytoplasmic puncta to regulate the availability of mDia1, thereby regulating its activity, which prevents the FH2 motif-mediated repression of MyoG expression due to high mDia1 activity. Although we have shown that mDia1 interacts with MyoD, pAkt2(Ser474), β-Catenin and Phb1, it remains unclear whether these interactors bind the cytoplasmic mDia1-Phb2 complex to regulate MyoG expression or exist as separate mDia1-interacting pools. Dotted box in MT indicates a possible mDia1 complex that might be involved in regulation of MyoG in MT, but needs additional studies. Dotted arrows represent mechanisms that have not been studied in MT.

## Discussion

We report that signalling mediated by the RhoA effector mDia1 during differentiation is anti-myogenic, and identify a new mDia1-interacting protein, the multi-functional Phb2 that mitigates these effects to facilitate progression of the myogenic program. We map the domains by which Phb2 and mDia1 interact. We demonstrate that mDia1 represses MyoG expression in MT, and that this repression is relieved by interaction with Phb2. We further demonstrate that mDia1 interacts with differentiation-promoting transcription factors MyoD, pAkt2(ser474) and active β-Catenin in MT. We report that mDia1 acts as a scaffold molecule with the potential to bind many proteins that might regulate its activity and impact multiple pathways in a stage-specific manner. Finally, we propose a model wherein mDia1 activity is fine-tuned by Phb2-mediated sequestration of mDia1 in cytoplasmic puncta in MT to promote MyoG expression during differentiation.

### mDia1 represses MyoD and MyoG expression in MT

Our studies place the RhoA effector mDia1 as a negative regulator of MyoD and MyoG expression during differentiation. Consistent with the repressive effects of RhoA on differentiation (Beqaj et al., 2002; Castellani et al., 2006; Charrasse et al., 2006; Gallo et al., 1999; Meriane et al., 2000; Nishiyama et al., 2004), we report that endogenous mDia1 suppresses MyoD and MyoG transcript expression. Similarly, mDia1ΔN3 suppressed MyoD transcription, and although it did not alter MyoD protein levels, it represses MyoD function. Interestingly, MyoG promoter activity, mRNA and protein expression were all reduced by ectopic constitutively active mDia1ΔN3, indicating that high mDia1 activity is repressive. Our findings are consistent with reports of MyoD-independent regulation of MyoG expression (Takano et al., 1998; Wilson and Rotwein, 2006), wherein ectopic expression of Rho-GDI or inhibition of IGF-II, repressed MyoG expression but did not affect MyoD. Taken together, our findings suggest that high mDia1 activity is anti-myogenic during differentiation, and may channel the known repressive effects of RhoA in myogenesis.

### Domain-specific interaction in MT permits Phb2 to relieve mDia1-mediated repression of MyoD and MyoG

Domain-specific interactions with proteins particularly in MT might have evolved to mitigate the repressive effects of mDia1 on differentiation, while protecting/permitting the common functions of mDia1 in both states. In this context, we show that the binding of Phb2 to mDia1 specifically in MT is pro-myogenic, counteracting the anti-myogenic effects of mDia1. Our findings contrast with the anti-myogenic role ascribed to Phb2 in previous studies (Héron-Milhavet et al., 2008; Sun et al., 2004), possibly since those reports used synthetic reporters containing control elements that do not reflect the endogenous promoters, and lacked direct loss-of-function studies in MT. In addition, those studies did not report the interaction of Phb1, a known interactor for Phb2 (Merkwirth and Langer, 2009; Nijtmans et al., 2000; Tatsuta et al., 2005), whereas we detect interactions of mDia1 with Phb2 along with Phb1, MyoD, active β-Catenin and phospho-Akt2.

Phb2 is a highly conserved and ubiquitously expressed protein (Bavelloni et al., 2015a; Mishra et al., 2006), that shows cell type-specific localisation to lipid rafts (Sharma and Qadri, 2004), mitochondria (Tatsuta et al., 2005), cytoplasm (Takata et al., 2007) or nucleus (Bavelloni et al., 2015b; Kasashima et al., 2006; Peng et al., 2015; Thuaud et al., 2013). While the molecular basis for its diverse subcellular locations is unknown, Phb2 clearly shows pleiotropic functions and is implicated in cell survival (Chowdhury et al., 2014; Peng et al., 2015; Thuaud et al., 2013), cell signalling (Bavelloni et al., 2015a), stem cell proliferation (Kowno et al., 2014) and gene regulation (Mishra et al., 2006). Consistent with reports of shuttling between mitochondria and the nucleus (Kasashima et al., 2006; Kuramori et al., 2009), and its role in these organelles, we show that Phb2 localises to both mitochondria and the nucleus in muscle cells. In the nucleus, Phb2 represses ERα-mediated transcription (Delage-Mourroux et al., 2000; Kurtev et al., 2004; Montano et al., 1999), and muscle-specific gene expression (Héron-Milhavet et al., 2008; Sun et al., 2004). Our findings add to Phb2’s transcriptional control function, as its interaction with mDia1 is pro-myogenic, wherein binding of Phb2 relieves the mDia1-mediated repression of MyoD and MyoG.

The domain mapping of Phb2-mDia1 interaction reveals mechanistic avenues. Phb2 binds to aa 752-978 region of mDia1, a region which we earlier reported to repress MyoD expression in MB (Gopinath et al., 2007). Further, the FH2 motif (aa 946-1010), which partially overlaps with Phb2 binding region on mDia1, also represses MyoG promoter activity. Conceivably, the binding of Phb2 to aa 752-978 region blocks the negative regulation of MyoG by mDia1. Additionally, upon binding to mDia1, Phb2 might recruit pro-myogenic regulators to the FH2 motif, promoting MyoG expression. The failure of the mDia1ΔN3(HindIII)-Phb2 interaction to relieve repression of MyoG promoter indicates that in the absence of recruitment of pro-myogenic regulators to the FH2 scaffold/FH2 motif, mDia1ΔN3(HindIII) functions as a dominant negative, sequestering Phb2 from its de-repressive role. In support of this hypothesis, we find that mDia1 binds pro-myogenic proteins MyoD, Akt2, and β-Catenin in MT. MyoD is required for MyoG transcriptional induction (Berkes and Tapscott, 2005b), Akt2 induces MyoG expression (Sumitani et al., 2002), while β-Catenin, a key effector of Wnt signaling promotes MyoD expression and function (Kim et al., 2008; Petropoulos and Skerjanc, 2002; Ridgeway et al., 2000; Suzuki et al., 2015). In addition, we show that the mDia1-associated Akt2 is phosphorylated at Ser474 (Tsuchiya et al., 2014), suggesting that mDia1 interacts with activated Akt2. Moreover, expression of Akt2 but not Akt1 correlated with differentiation (Calera and Pilch, 1998; Gonzalez et al., 2004; Héron-Milhavet et al., 2006; Héron-Milhavet et al., 2008; Vandromme et al., 2001). Further, Insulin and Wnt pathways cooperate to promote myogenesis (Rochat et al., 2004). Currently, it is unclear which of these proteins is responsible for rescue of MyoG expression. Our studies identify mDia1 as a node for mediating interactions with several proteins, which in turn might mitigate its anti-myogenic functions in MT.

Earlier studies have reported that activity of mDia1 and its isoforms needs to be tightly regulated to promote optimal function (DeWard and Alberts, 2009; Gopinath et al., 2007; Li and Sewer, 2010). Post-translational modifications such as ubiquitination (DeWard and Alberts, 2009), phosphorylation and sumoylation (Li et al., 2013) have been reported to regulate mDia stability and function. Our studies also provide evidence that mDia1 activity needs to be dampened in order to promote differentiation, and the reduction of mDia1 expression during myotube formation supports this notion (Fig.S3A). We detected a cytoplasmic interaction of mDia1 with Phb2, wherein mDia1 and Phb2 co-localised in cytoplasmic puncta in MT. This finding suggests that Phb2 sequesters mDia1 in these puncta thereby regulating its activity and restricting nuclear entry. Our findings are consistent with the studies that report that RhoA activity needs to be down regulated prior to myoblast fusion to promote differentiation (Charrasse et al., 2006; Doherty et al., 2011; Fortier et al., 2008; Nishiyama et al., 2004) and provide a mechanism for down regulating RhoA activity to promote diffentiation that controls the availability of mDia1 downstream RhoA, through cytoplasmic sequestration by Phb2, thereby preventing its anti-myogenic functions. It can be speculated that upon binding Phb2, mDia1 switches from anti-myogenic to pro-myogenic, acting as a scaffold to form a pro-myogenic complex with MyoD, pAkt2(Ser474), β-Catenin or Phb1 during differentiation. Although Phb1 has not been reported to regulate myogenesis, it represses E2F dependent transcription (Wang et al., 1999a; Wang et al., 1999b; Wang et al., 2002) and might promote irreversible cycle exit during differentiation. Taken together, these findings suggest the formation of a pro-myogenic complex restricting stage-specific mDia1 functions during differentiation. Our data currently do not allow us to distinguish whether pAkt2, MyoD, Phb1 and active β-Catenin simultaneously associate with the mDia1-Phb2 complex or exist as different mDia1-bound complexes. In conclusion, we propose a model where Phb2 sequesters mDia1 in the cytoplasm of MT to regulate its anti-myogenic activity thereby preventing the mDia1-mediated repression of MyoG, the nodal transcription factor required for orchestrating the myogenic cascade.

### Analysis of mDia1-interacting proteins reveals stage-specific interaction with regulators of cytoskeletal dynamics, signaling, metabolic functions and myogenesis

LC-MS/MS analysis of mDia1-interacting proteins revealed that mDia1 acts as a scaffold with both common and stage-specific partners which govern its functions in MB and MT, and that its indispensable role in cytoskeletal dynamics common to both states is well preserved. mDia1 associates with cytoskeletal regulators in both MB and MT, where its role in microfilament dynamics is preserved. We confirm that mDia1 associates with the actin binding proteins (ABP) profilin, a known interactor (Watanabe et al., 1997) that controls polymerisation, and cofilin1, that regulates actin disassembly (Ghosh, 2004) in both MB and MT. Additionally, the role of mDia1 in mediating cytoskeletal signalling through GTPases is preserved in both MB and MT, although the associated GTPases are different. Association of mDia1 with exosomal and vesicle proteins in both MB and MT, highlights another cytoskeleton-dependent mDia1 function that is maintained in these stages and is consistent with the previously reported roles for Rho in vesicle trafficking (Symons and Rusk, 2003), endosome dynamics (Ellis and Mellor, 2000; Gasman et al., 2003a; Sandilands et al., 2004), exocytosis (Gasman et al., 2003b). RhoA-GTPase and actin dynamics are established regulators of vesicle trafficking in both endocytic and exocytic pathways (Ridley, 2001; Ridley, 2006). Exosomes are involved in directed migration of cells in tissues (Sung et al., 2015). Therefore, mDia1 might coordinate cytoskeletal changes with vesicle dynamics to regulate exosome-specific roles in these states. Taken together, we report that the cytoskeletal function of mDia1, which is critical in regulating various cellular processes is maintained in both MB and MT.

Unlike in MB, mDia1 binds a plethora of proteins belonging to distinct classes such as metabolic, mitochondrial, and proteasomal in MT, reflecting its significance in regulating differentiation. MT are highly metabolically active (Leary et al., 1998; Wagatsuma and Sakuma, 2013) and the association of mDia1 with a multitude of metabolic proteins suggests a role for mDia1 in these metabolic processes and differentiation. Association of mDia1 with mitochondrial proteins in our studies is consistent with its reported role in regulating mitochondrial trafficking in adrenocortical cells (Li and Sewer, 2010). Interestingly, mitochondria have been reported as potential regulators of myogenesis, wherein mitochondrial respiration and enzyme activity increase during differentiation, while perturbation of mitochondrial activity blocks differentiation (Wagatsuma and Sakuma, 2013). Mitochondria regulate insulin-mediated myogenesis through c-Myc and Calcineurin (Friday et al., 2003; Pawlikowska et al., 2006; Seyer et al., 2006; Seyer et al., 2011). MyoG but not MyoD expression is directly induced by mitochondrial activity in avian MB (Rochard et al., 2000). This is consistent with the observed MyoD-independent regulation of MyoG, which suggests that mitochondria may regulate MyoG expression in MT, downstream of mDia1-Phb2 interaction. Additionally, Phb2 regulates mitophagy (Wei et al., 2017) and mitophagy is required for myoblast differentiation (Sin et al., 2016), suggesting a possible role for mDia1 in regulating mitophagy and differentiation. Our findings suggest that mDia1 might regulate metabolic processes, mitochondrial physiology and mitophagy to regulate differentiation. Interestingly, we report that mDia1 associated with several proteasomal proteins, similar to mDia2, an isoform of mDia1 (Isogai et al., 2015), but it remains to be understood whether mDia1 is being targetted or targets other proteins for degradation. Nonetheless, ubiquitination and proteasomes have been reported to regulate RhoA-GTPase activity (de la Vega et al., 2011; Doye et al., 2002) and it can be speculated that mDia1 might be targetted by the proteasome to regulate its activity. However, it is not unlikely, that mDia1 itself might target other proteins for turnover through its association with proteasomal proteins. Interestingly, the proteasome is also required for mitochondrial protein quality-control and health, suggesting that mDia1 might regulate these processes as well (Bohovych et al., 2015; Radke et al., 2008). Our study reports novel stage-specific roles for mDia1 in MT and suggests that mDia1 modulates mitochondrial, metabolic and proteasomal functions to regulate differentiation.

The stage-specific role of mDia1 in regulating muscle-specific gene expression is strengthened by the muscle-specific proteins that we found as partners in MB and MT. These include, Galectin3 (Rancourt et al., 2017), Reticulon-4 (Magnusson et al., 2003) and Calponin (Duband et al., 1993; Michael et al., 1992) in MT, and Annexin A1 in both MB and MT (Bizzarro et al., 2012; Leikina et al., 2015). Association of mDia1 with proteins controlling adhesion such as Vinculin, Talin (Humphries et al., 2007), Pdlim1, (Chen et al., 2016) and Ras suppressor protein 1 (Dougherty et al., 2005) in MT, suggests a role for mDia1 in myoblast fusion during differentiation as reported in flies (Deng et al., 2015; Deng et al., 2016). mDia1 plays an antagonistic role in regulating muscle-specific gene expression: while our previous studies showed ectopic mDia1 repressed MyoD protein levels in MB (Gopinath et al., 2007), it did not affect MyoD protein levels in MT (this study). Additionally the knockdown of endogenous mDia1 in MB reduced MyoD expression (Gopinath et al., 2007) while, knockdown of mDia1 in MT induced MyoD expression in our study, again pointing to stage-specific roles. We report that mDia1 is anti-myogenic, as indicated by knockdown and over-expression studies. Interestingly, the binding of Phb2 to mDia1 in MT alleviates its repressive effects on muscle-specific gene expression, thereby affecting only one of the several functions of mDia1 uncovered in our studies. We propose that mDia1 may play antagonistic roles in MB and MT: while its activity in MT is balanced by interacting proteins that mitigate its anti-myogenic functions and promote its pro-myogenic functions, its common functions in both MB and MT may be preserved as they are essential for cytoskeletal dynamics.

mDia1 interacts with proteins involved in nuclear import and protein folding in MB, suggesting novel stage-specific roles for mDia1 in MB. It’s interaction with proteins regulating nuclear import such as Kpnb1, Hsp90ab1and Hsp90aa1 (Hasse and Fitze, 2016; Stelma and Leaner, 2017; Zhong et al., 2014) suggests that mDia1 might shuttle between the nucleus and cytoplasm, similar to mDia2 (Miki et al., 2009; Shao et al., 2015). Consistent with its scaffolding properties (Wallar and Alberts, 2003), mDia1 associates with chaperones such as Hsp90b1, Hsp90ab1, Hsp90aa1 (Schopf et al., 2017), suggesting a potential involvement in regulating protein folding. Overall, our study reports novel stage-specific roles for mDia1 in MB and identifies several interactors which might help understand its diverse functions.

In conclusion, we suggest that mDia1 expression is retained in MT, despite its anti-myogenic effects, since it plays an indispensable role in cytoskeletal dynamics. The lower levels of mDia1 in MT might facilitate in maintaining a moderate level of RhoA signalling to prevent anti-myogenic effects of hyper-active RhoA-mDia1 signalling. We show that mDia1 has stage-specific roles in MB and MT and that these roles are modulated by stage-specific interactions with proteins that mitigate only those functions of mDia1 that are deleterious to that stage. We identified Phb2 as one such mDia1-interacting protein in MT that regulates the activity of mDia1 by controlling the availability of mDia1 downstream RhoA through sequestration, since hyper-active mDia1 activity is anti-myogenic. Thus, Phb2 functions to maintain a pro-myogenic level of mDia1 activity and RhoA signalling to promote MyoG expression and differentiation. As a result, mDia1 when bound to Phb2 switches its role from anti-myogenic to pro-myogenic. Taken together, we report that mDia1 in the absence of Phb2 interaction is anti-myogenic and might be involved in suppressing MyoG expression in MB, while in MT, owing to its reduced expression and regulated activity due to its interaction with Phb2, is pro-myogenic and might promote differentiation by regulating mitochondrial, metabolic and proteasomal functions.

## Acknowledgements

We thank our colleagues Suchitra Gopinath (THSTI) and Ghanshyam Swarup (CCMB) for critical comments on the manuscript. This work is part of the doctoral thesis of AS (Manipal Academy of Higher Education) who was supported by a graduate fellowship from the Council of Scientific and Industrial Research (CSIR), and core funds from CCMB. We also acknowledge a Govt. of India Department of Science and Technology, SERB National Postdoctoral Fellowship to GS, core support from DBT to InStem, core funds from CSIR to CCMB, and grants from the Indo-Australia Biotechnology Fund (DBT) and the Indo-Danish Strategic Research Fund (DBT) to JD. We gratefully acknowledge the Proteomics facility at CSIR-CCMB, flow cytometry and imaging facilities (CIFF) at NCBS-InStem and Advanced Imaging Facility at CSIR-CCMB.

## Materials and Methods

### Yeast two hybrid screen

*Saccharomyces cerevisiae* strain PJ69-4A was co transformed with mDia1ΔN3-BD (pGBKT7–GAL4 Binding domain vector) and Matchmaker 7day old mouse embryonic cDNA library cloned in pACT2 (AD-GAL4 Activation domain vector) (Clonetech) as per manufacturer’s instructions. Briefly, transformants were plated on amino acid dropout selection plates lacking Trp (tryptophan), Leu (leucine) and Adenine (Ade). Reporters *ADE2* encoding Ade biosynthesis enzymes and *LacZ* encoding β-galactosidase were used to identify putative mDia1-interacting proteins. Reporter expression was assessed by plating on selection plates – TLA (-Trp/-Leu/-Ade) for screening *ADE2* expression indicated by growth and – TL+X-Gal (-Trp/-Leu/+X-Gal) for LacZ expression indicated by blue pigmentation. PJ69-4A cotransformed with *Drosophila* Trithorax and GAGA factor were used as a positive control whereas co-transformation with empty pGBKT7 and pACT2 vectors served as a negative control. Clones positive for expression of both reporters were selected and re-screened three times serially for reporter expression. The AD plasmid was isolated from positive clones derived from single yeast colonies, screened for the presence of insert after transforming yeast DNA into E.coli DH5α and subjected to hybrid reconstitution assays. For reconstitution, PJ69-4A was co-transformed with AD vector from the positive clone and mDia1ΔN3-BD or empty pGBKT7, followed by plating on –TLA and –TL+X-Gal plates. Four colonies for each co-transformation per clone were screened three times serially on reporter plates. The AD plasmid from yeast clones that remained positive throughout was then isolated, transformed into E.coli DH5α and sequenced. Identity of the sequenced clones was determined by performing NCBI-nucleotide BLAST analysis against mouse genomic Reference RNA (Ref seq_RNA) database.

### Cell culture

Mouse C2C12 subclone A2 MB (C2C12 obtained from Helen Blau, Stanford, subcloned in the lab (Sachidanandan et al., 2002) were cultured under proliferative condtions using growth medium (GM; DMEM+20% FBS). MB were differentiated into MT by culture in low serum differentiation medium (DM; DMEM+2% Horse serum) for 24, 36, 72 or 120 hours, replenished daily. HEK293T cells were cultured in DMEM +10% FBS. All media were supplemented 100 units/ml Penicillin, 100μg/ml Streptomycin (Cat. no. 15140-163, Thermo scientific) and 2 mM Glutamax (Cat. no. 35050-079, Thermo Scientific). DMEM (Cat. no. 10313-021, Thermo Scientific), FBS (Cat. no. 16000-044, Thermo Scientific) and Horse serum (Cat. no. 16050122, Thermo Scientific).

### Transfections

C2C12 MB or HEK293T were plated on coverslips or tissue culture dishes 12-16 hours prior to transfection using Lipofectamine LTX with plus reagent (Cat. no. - 15338-100, Thermo scientific), as per manufacturer’s instructions. 12 hours post transfection, GM was replaced by DM for 36 or 72 hours. Transfected cells were processed either for immunostaining, RNA extraction or dual Luciferase assay. Normalised DNA amounts were used for transfection to get similar expression levels of all mutants used in the dual luciferase assay. HEK293T cells were transfected 16 hours post plating using Lipofectamine LTX with plus reagent as per manufacturer’s instructions for 24 hours, followed by lysate preparation for western blot analysis or immunoprecipitation. For siRNA studies, MB were plated 16 hours prior to transfection with siGenome SMART pool siRNA from Dharmacon using RNAiMax (Cat.no.-13778-150, Invitrogen) as per manufacturer’s instructions. 12 hours post transfection, MB were trypsinised and plated for 48 hours in differentiation medium. Knockdown cells were harvested and processed for RNA extraction and qRT-PCR analysis. siRNA used in the study are mDia1 (Cat. no. M-064854-02-0050, Dharmacon), Phb2 (Cat. no. M-040938-01-0005, Dharmacon) and scrambled (SCR) control (Cat. no. D-001206-14-20, Dharmacon).

### Plasmids and Cloning

Expression plasmids for GFP-tagged mouse mDia1, mDia1FL, mDia1ΔN3, mDia1F2, mDia1ΔN3(HindIII), mDia1H+P and mDia1CC were gifts from S Narumiya (Watanabe et al., 1999). mDia1ΔN3-BD was cloned from mDia1ΔN3-pET28a into pGBKT7 (GAL4 binding domain vector-Clonetech) using Nde1 and BamH1. Flag-tagged mouse Phb2 expression plasmid Phb2-Y2H (89-299aa) was cloned into pCMV2B from pACT2 using EcoR1 and Xho1. Flag tagged mouse Phb2 truncated mutants, Phb2 amino (89-180aa), Phb2 central (140-244 aa), Phb2 carboxy (180-299 aa) and Phb2 120-232 (120-232 aa) were cloned from Phb2-Y2H into pCMV2B using BamH1 and Xho1. Flag-tagged mouse Phb2-FL was obtained from Origene. MyoG prom-pGL3 was a gift from Eric Olson’s lab (Edmondson et al., 1992b). pRLSV40 Renilla Luciferase plasmid and pBluescript KS were obtained from Addgene.

### RNA isolation and analysis

MB were transfected with over-expresson constructs or siRNA for 12 hours in GM, followed by addition of DM for 36 or 48 hours. Cells were washed with ice cold PBS twice followed by lysis with Trizol (Cat no. 15596-026, Thermo Scientific), from which RNA was isolated as per manufacturer’s instructions, & purified by treatment with DNAase (Cat no. AM1906, Ambion). 1 μg of total RNA was used to synthesize cDNA using Superscript III (Cat no. 18080-044, Thermo Scientific) and amplified by qRT-PCR (master mix was made with cDNA diluted 1:5, primers and Maxima SYBR Green 2X PCR master mix (Cat no.K0222, Fermentas) and analysed in triplicate on a ABI 7900HT thermal cycler (Applied Biosystems). Amplicons were verified by sequencing and dissociation curves. Relative level of endogenous MyoD and MyoG mRNA in the transfected samples was calculated with respect to untransfected sample or SCR control after normalising to corresponding GAPDH levels in the transfected samples. Fold change between samples was calculated using [2^(–ΔΔCt)^] method. Primers used in the study:, GAPDH 5’-AAGGCCGGGGCCCACTTGAA-3’, 5’-AGCAGTTGGTGGTGCAGGATGC-3’; MyoD 5’-ATGGCATGATGGATTACAGCGGCC-3’, 5’-GCTCCACTATGCTGGACAGGCAG-3’; MyoG 5’-CAACCAGCGGCTGCCTAAAGTGG 3’, 5’-GCATTCACTGGGCACCATGGGC-3’.

### Immunostaining

MB plated on coverslips were cultured in either GM or DM for 72h (D72), followed by fixation with 4% PFA in PBS at room temperature (RT) for 15 min. For transfected MT, growth medium was replaced 12 hours after transfection with DM for 36 hours. MT were fixed with 4% PFA in PBS for 15 min at RT. For immunostaining, cells were permeabilised with PBS+0.5% Triton-X-100 for 1 hour, followed by blocking with PBS+0.25% Triton-X-100+10% FBS for 1 hour at RT. Primary antibody incubations were performed overnight at 4°C, followed by three washes at RT with PBS+0.025% Tween 20, and detection with Alexa-fluor conjugated secondary antibodies for 1 hour prior to staining with DAPI (1μg/ml) for 10 min nuclear to reveal nuclei and mounting in Fluormount (Cat no. 0100-01, Southern Biotech). Confocal images were acquired on a confocal laser scanning microscope (Leica TCS SP8, Germany) using HC PL APO CS2 40X/1.3 Oil immersion objective at Zoom 1.28 for over-expression studies and HC PL APO CS2 63X/1.4 Oil immersion objective at Zoom 3 for mDia1-Phb2 colocalisation or Zoom 2 for Organelle staining.

### Western blot analysis

Whole cell lysates were prepared in 2X SDS lysis buffer (100mM Tris-HCl pH 6.8, 4% SDS, 10% β-mercaptoethanol and 10mM EDTA) and 20-40 μg of whole cell lysate or equal volume of IP product was separated by SDS-PAGE followed by transfer to Polyvinylidene Difluoride (PVDF) membrane (Cat. no. 162-0177, Biorad). Primary antibody incubation was for 1 hour at RT or overnight at 4°C, followed by incubation with HRP-conjugated secondary antibody for 1 hour at RT. Chemiluminescent signal was detected by ImageQuant (Amersham) or ChemDoc (Syngene) using ECL detection reagent (Amersham). Antibody dilutions are listed in Table S7.

### Immunoprecipitation assays

Cells were washed once with cold PBS, lysed in modified RIPA buffer (50 mM Tris-HCl pH 7.4, 150 mM NaCl, 1% NP40, 0.25% Sodium deoxycholate and 1 mM EDTA) or mDia1 IP buffer (10 mM Tris-HCl pH 7.5, 150 mM NaCl, 1 mM EDTA, 1 mM EGTA, 10% Sucrose and 1% TX100) containing 1X protease and phosphatase inhibitors for 0.5-2h at 4°C, and cleared by centifugation at 13,000 rpm at 4°C for 20 min. Prior to pulldown, lysates containing equal protein were pre-cleared using BSA (10 μg/ml) and Protein A or G agarose beads for 1 hour at 4°C, then incubated with 2-3 μg primary antibody against flag, mDia1 or Phb2 for 16 hours at 4°C, followed by addition of Protein A or G agarose beads for 8 hours at 4°C. Immune complexes were collected by centrifugation at 2000 rpm for 10 min at 4°C, followed by three washes with cold PBS+0.5% Triton-X-100. The agarose beads were boiled in equal volume of Laemmli sample buffer for 5 min at 95°C to elute the bound immuno-precipitated proteins from the beads and IP eluates were collected after centrifuging at 13,000 rpm for 5 min at RT. Equal volume of IP product was subjected to western blot analysis.

### Cytoplasmic and nuclear fractionation

D72 MT were trypsinised, washed twice with cold 1X PBS and resuspended in 10 times the pellet size of Dia lysis buffer containing 0.2% TX100, 1X protease inhibitors and phosphatase inhibitors. Samples were incubated on ice for 10 min, followed by gentle vortexing for 15 sec. Cytoplasmic fraction was collected after two serial centrifugations at 4°C for 15 min at 800g. The nuclear pellet was washed four times with F2 buffer without detergent (20 mM Tris-HCl pH 7.6, 0.1 mM EDTA and 2 mM MgCl_2_) and lysed in mDia1 IP buffer containing 1X protease inhibitors and phosphatase inhibitors for 2 hours at 4°C. Nuclear fractions were collected by centrifugation at 13,000 rpm for 20 min at 4°C. Cytoplasmic and nuclear fractions were used for immunoprecipitation studies.

### Dual-Luciferase assays

MB were plated in 24 well dishes in triplicate 16 hours prior to transfection with pRLSV40 Renilla Luciferase (Addgene), MyoG-promoter/empty pGL3 Luciferase reporter constructs, mDia1ΔN3/ mDia1H+P, Phb2-FL/ Phb2 carboxy/Phb2 amino, empty pEGFPC1 or empty pCMV2B constructs. pBluescript KS (pBSKS) was used to ensure equal DNA amount during transfection. 12 hours post transfection, cells were shifted to DM for 72 hours. Reporter gene expression was assayed using Dual-Luciferase kit as per manufacturer’s instructions (Cat. no. E1910, Promega). Briefly, cells were lysed in the 1X PLB buffer, followed by addition of luciferase assay reagen LARII to record firefly Luciferase activity in a TD-20/20 luminometer (Turner Designs). Stop and Glow was then added to record Renilla Luciferase activity. Luciferase readings were expressed as relative light units (RLU) normalised to Renilla Luciferase for transfection and pGL3 for basal Luciferase activity.

### Mass spectrometric analysis

mDia1 was immunoprecipitated from lysates from GM and D72 cultures and IP confirmed by western blotting, following which IP products from GM and D72 were loaded onto a NuPAGE 4-12% Bis-Tris pre-cast gradient gels (Invitrogen). Electrophoresis was performed at 200 V using MES running buffer for approximately 40 min. Proteins were visualize by staining with Coomassie brilliant blue R250 and each Coomassie stained lane was processed individually by division into 4-5 pieces containing approximately 2-3 bands. Each of these pieces was individually cut into smaller pieces (1-2mm), in-gel digested, desalted and enriched for Liquid chromatography tandem mass spectrometry (LC-MS/MS) analysis as described (Rappsilber et al., 2003; Shevchenko et al., 1996). Briefly, eluted peptides from desalting tips were resuspended in 2% (v/v) formic acid and sonicated for 5 min. Samples were analyzed on Q Exactive Hybrid Quadrupole-Orbitrap Mass spectrometer (Thermo Scientific) coupled to a nanoflow LC system (Easy nLC II, Thermoscientific). Peptide fractions were loaded onto a BioBasic C18 PicoFrit 15 μm nanocapillary reverse phase HPLC column (75 μm × 10 cm; New Objective, MA, USA) and separated using a 60 min linear gradient of the organic mobile phase [5% Acetonitrile (ACN) containing 0.1% formic acid and 95% ACN containing 0.1% formic acid], at a flow rate of 400 nl min^−1^. Q Exactive Hybrid Quadrupole-Orbitrap Mass spectrometer (Thermo Scientific) was used for the analysis. Protein/peptides were identified by searching against UniProt/Swissprot amino acid sequence database of *Mus musculus* (release March 2016 with 16790 entries) and a database of known contaminants using MaxQuant software (Version 1.3.0.5) (Cox and Mann, 2008). MaxQuant uses a decoy version of the specified UniProt database to adjust the false discovery rate for proteins and peptides below 1%. The search was set up for tryptic peptides with minimum peptide length of seven aa, including constant modification of cysteine by carbamidomethylation, minimum two peptide identification and label-free quantitation (LFQ). LFQ ratio for individual proteins was calculated by LFQ in mDia1 IP/LFQ in IgG. Proteins that had LFQ ratio of 2 or greater were selected for further analysis. Three independent biological samples of MB and MT were processed by mDia1 IP-LC-MS/MS and only those proteins that were detected with significance in all three runs were selected for further analysis. Gene ontology analysis was performed using REVIGO http://revigo.irb.hr.

---

**Figure S1.**
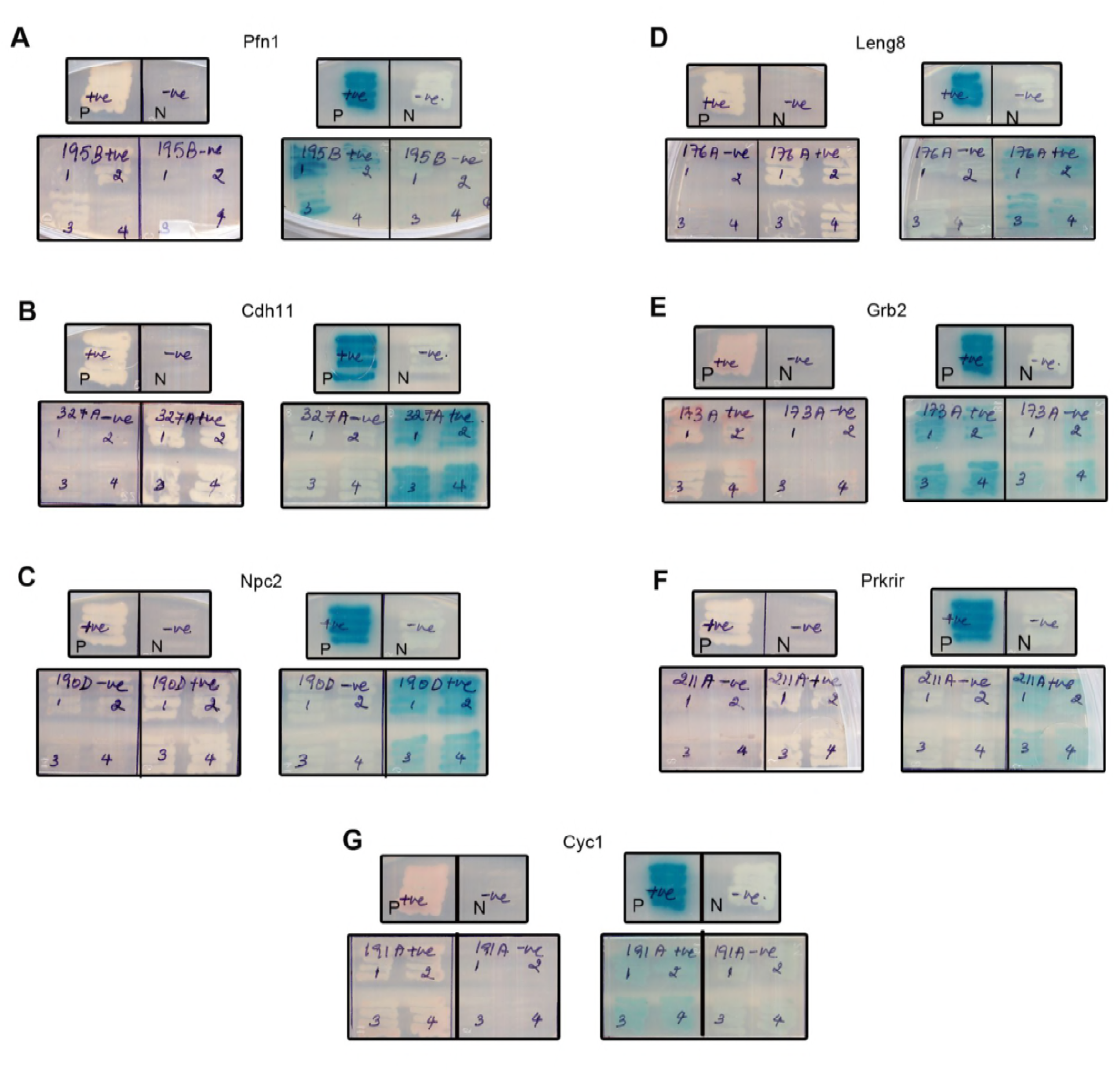
Putative mDia1-interacting proteins identified in the yeast two hybrid screen. Yeast two-hybrid screen to identify novel mDia1-interacting proteins. A GAL4 hybrid reconstitution assay with putative mDia1 interactors was performed to study the induction of reporters *ADE2* and *LacZ* on -Trp/-Leu/-Ade and -Trp/-Leu+X-Gal respectively. PJ69-4A was co transformed with interacting protein plasmid and mDia1ΔN3-BD (positive reconstitution) or empty BD vector (negative reconstitution). Four colonies per per reconstitution assay (positive and negative) were screened on selection plates. PJ69-4A co-transformed with *bona fide* interacting proteins *Drosophila* Batman-AD and GAGA factor-BD served as a positive control “P” for reporter expression and PJ69-4A co-transformed with empty pACT2 and pGBKT7 vectors served as a negative control “N”. Induction of *ADE2* reporter is indicated by growth and induction of *LacZ* expression is indicated by blue colour in the colonies on the selection plates. Seven library clones that induced the reporter expression were finally selected from the screen as mDia1-interacting proteins of interest. (A) Profilin1 (Pfn1), known interactor for mDia1 (B) Cadherin11 (Cdh11) (C) Niemann Pick Type C2 (Npc2) (D) Leukocyte receptor cluster (LRC) member 8 (Leng8) (E) Growth factor receptor bound protein 2 (Grb2) (F) Protein-kinase, interferon-inducible double stranded RNA dependent inhibitor, repressor of (p58 repressor) (Prkrir) (G) Cytochrome c1 (Cyc1) were identified as mDia1-interacting proteins. Trp-Tryptophan, Leu-Leucine, Ade-Adenine.

**Figure S2.**
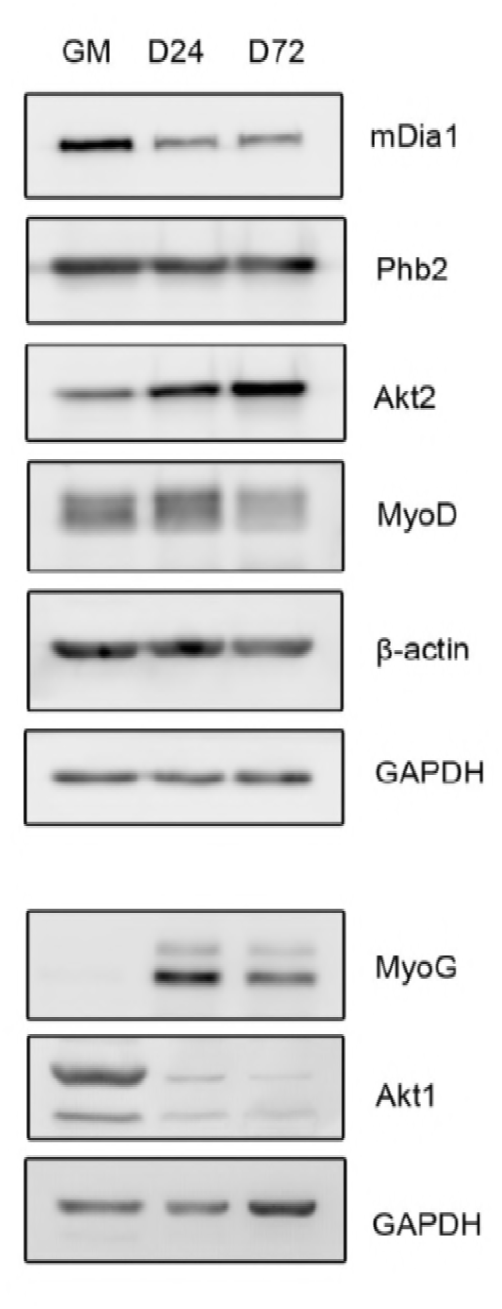
Expression profile of mDia1 and Phb2 during differentiation. Western blot to study expression profile of mDia1 and Phb2 during differentiation. Lysates were harvested from proliferating C2C12 (GM), differentiated C2C12 maintained in differentiation medium for 24h (D24) or 72h (D72) and subjected to western blotting using various antibodies. Left side of panel are the corresponding sizes in kDa.

**Figure S3.**
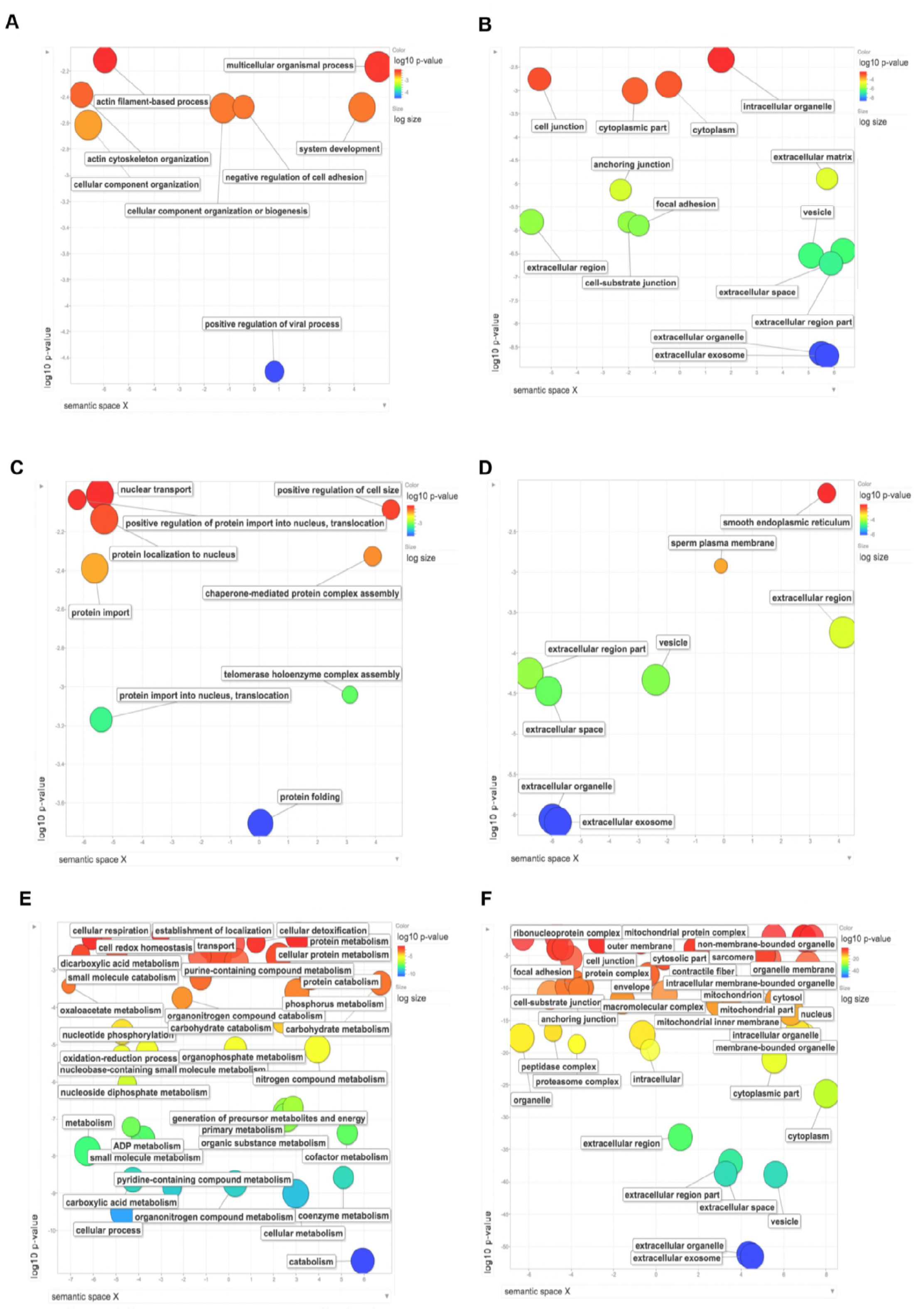
Gene Ontology of mDia1-interacting proteins identified in MB and MT by mDia1 IP-LC-MS/MS analysis. Gene ontology analysis of mDia1-interacting proteins identified by mDia1 IP-LC-MS/MS analysis in myoblasts (MB) and myotubes (MT) lysates was performed using REVIGO based on associated biological processes and cellular components. Gene ontology of proteins that bind mDia1 in both MB and MT based on (A) biological process and (B) cellular components. Gene ontology of mDia1-interacting proteins in MB based on (C) biological process and (D) cellular components. Gene ontology of the proteins that bind mDia1 in MT based on (E) biological process and (F) cellular components.

**Figure S4.**
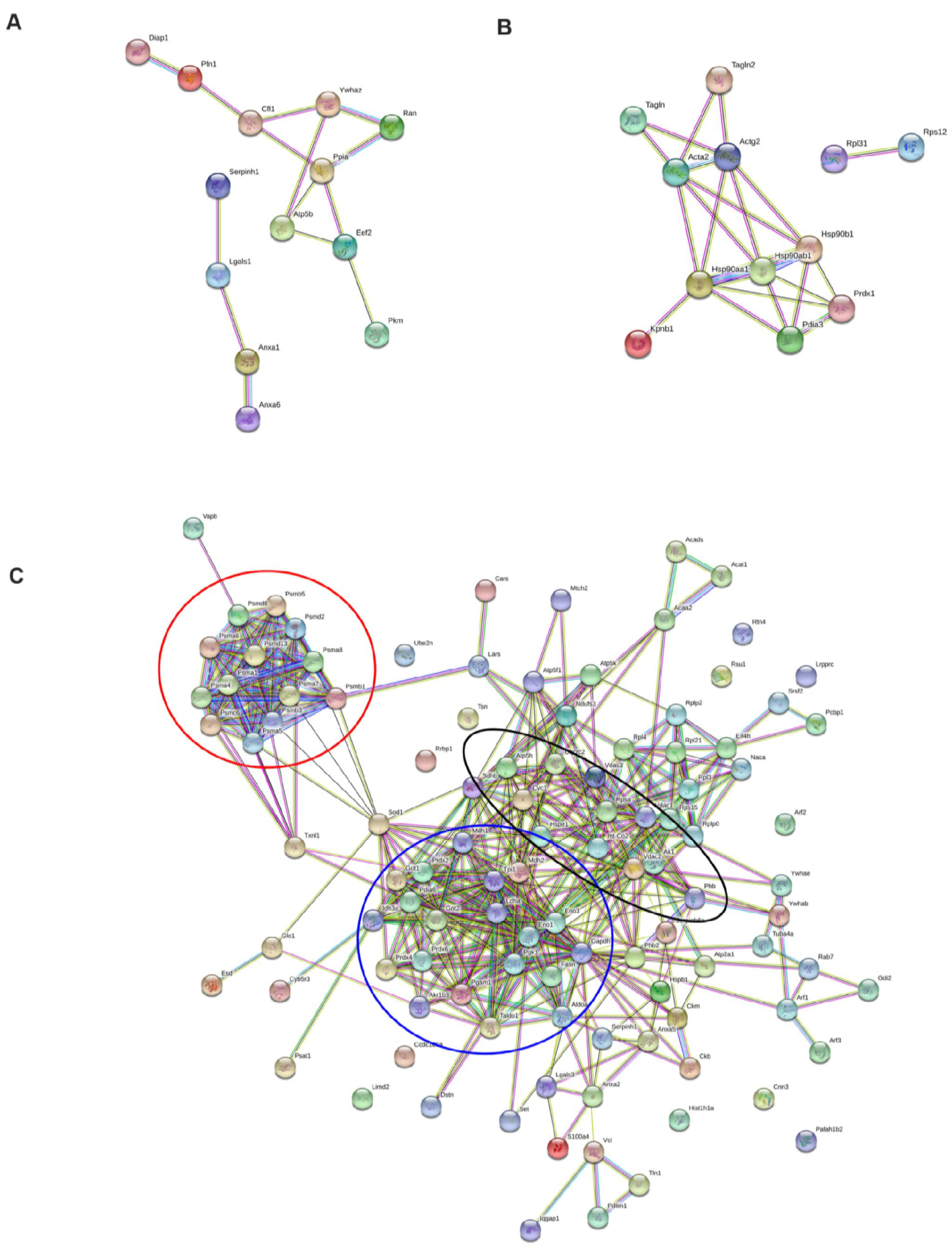
Protein association networks for mDia1-interacting proteins in MB and MT generated by STRING. STRING analysis of mDia1-interacting proteins in (A) both MB and MT, (B) MB and (C) MT. Highlighted clusters-Proteasomal proteins (Red), metabolic enzymes (Blue), mitochondrial proteins (Black).

**Figure S5.**
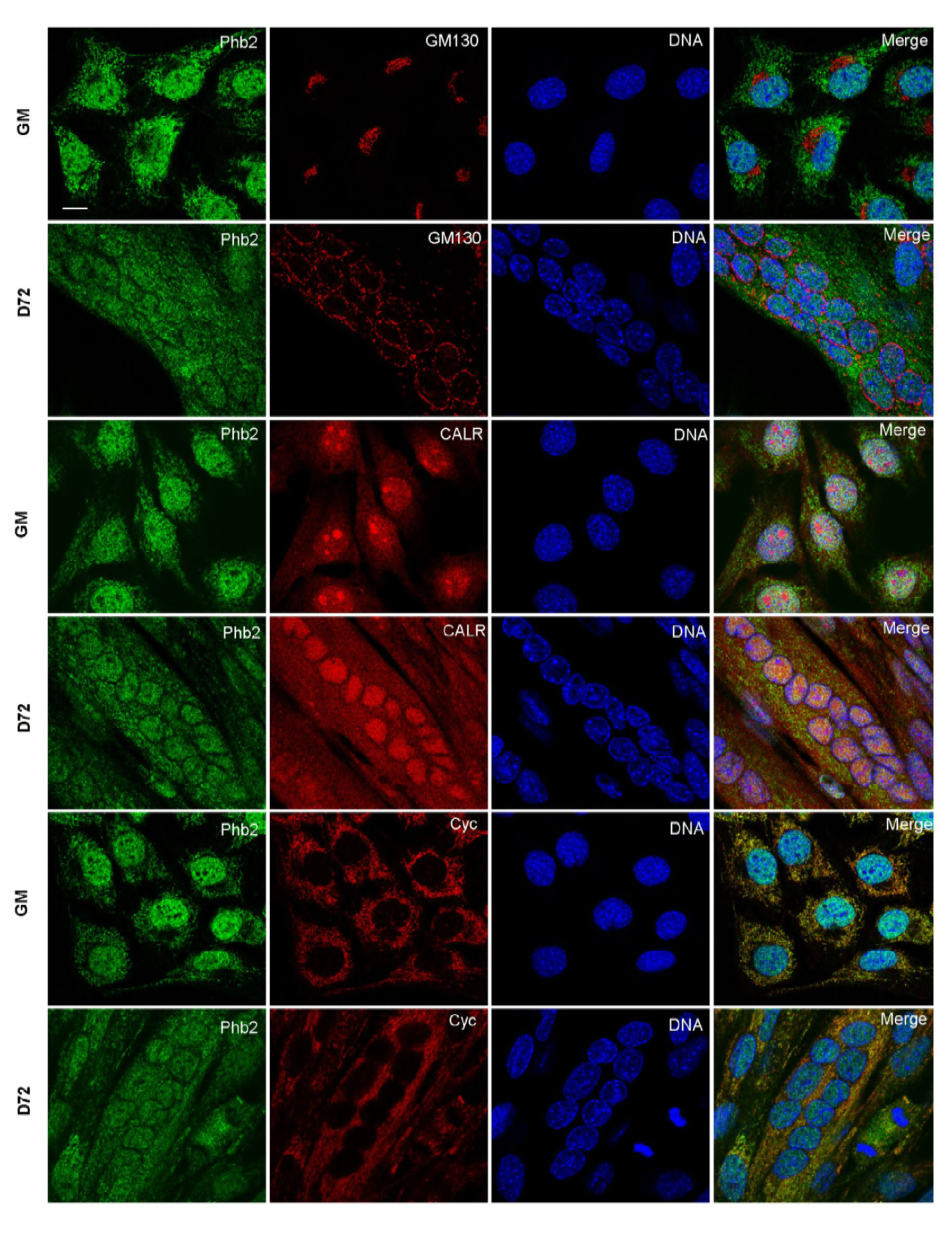
Localisation of Phb2 during differentiation. Cellular localisation of Phb2 during proliferation and differentiation of C2C12. MB (GM) and MT (D72) were fixed and co-immunostained by antibodies against Phb2 and organelle markers GM130, Cyc or CALR. (Cyc-Cytochrome c-Mitochondrial marker, GM130-cis-Golgi matrix protein-Golgi marker, CALR-Calreticulin-Endoplasmic reticulum marker). Confocal images were acquired using Leica TCS SP8 confocal microscope. Scale bar represents 10 μm.

**Table S1.** mDia1-interacting proteins identified in Y2H.

| S.no | Name | Symbol | Yeast clone | Gene ID |
| --- | --- | --- | --- | --- |
| 1 | 195B | Profilin 1 | Pfn1 | 18643 |
| 2 | 194A | Prohibitin2 | Phb2 | 12034 |
| 3 | 169A, 173A | Growth factor receptor bound protein 2 | Grb2 | 14784 |
| 4 | 190D, 214A | Niemann Pick type C2 | Npc2 | 67963 |
| 5 | 175A, 291A, 292A, 327A | Cadherin 11 | Cdh11 | 12552 |
| 6 | 176A | Leukocyte receptor cluster (LRC) member 8 | Leng8 | 232798 |
| 7 | 211A | Protein-kinase, interferon-inducible double stranded RNA dependent inhibitor, repressor of (P58 repressor) | Prkrir | 72981 |
| 8 | 191A | Cytochrome c-1 | Cyc1 | 66445 |

**Table S2:** mDia1-interacting proteins commonly identified in MB and MT by LC-MS/MS.

| S.no. | Name | Symbol | UniProt ID |
| --- | --- | --- | --- |
| 1 | Protein diaphanous homolog 1 | Diaph1 | O08808 |
| 2 | Annexin A1 | Anxa1 | P10107 |
| 3 | Annexin A6 | Anxa6 | P14824 |
| 4 | Cofilin-1 | Cfl1 | P18760 |
| 5 | Elongation factor 2 | Eef2 | P58252;O08810 |
| 6 | Galectin-1 | Lgals1 | P16045 |
| 7 | Profilin-1 | Pfn1 | P62962;CON__P02584 |
| 8 | Pyruvate kinase isozymes M1/M2 | Pkm2 | P52480;P53657 |
| 9 | Peptidyl-prolyl cis-trans isomerase A | Ppia | P17742 |
| 10 | GTP-binding nuclear protein Ran | Ran | P62827;Q61820 |
| 11 | Serpin H1 | Serpinh1 | P19324 |
| 12 | 14-3-3 protein zeta/delta | Ywhaz | P63101;P62259;O70456 |
| 13 | ATP synthase subunit beta, mitochondrial | Atp5b | P56480 |

**Table S3:** mDia1-interacting proteins identified specifically in MB by LC-MS/MS.

| S.no. | Name | Symbol | UniProt ID |
| --- | --- | --- | --- |
| 1 | Actin, aortic smooth muscle;Actin, gamma-enteric smooth muscle | Acta2;Actg2 | P62737;P63268 |
| 2 | Heat shock protein HSP 90-alpha | Hsp90aa1 | P07901 |
| 3 | Heat shock protein HSP 90-beta | Hsp90ab1 | P11499 |
| 4 | Endoplasmin | Hsp90b1 | P08113 |
| 5 | Importin subunit beta-1 | Kpnb1 | P70168 |
| 6 | Protein disulfide-isomerase A3 | Pdia3 | P27773 |
| 7 | Peroxiredoxin-1 | Prdx1 | P35700 |
| 8 | 60S ribosomal protein L31 | Rpl31 | P62900 |
| 9 | 40S ribosomal protein S12 | Rps12 | P63323 |
| 10 | Transgelin | Tagln | P37804 |
| 11 | Transgelin-2 | Tagln2 | Q9WVA4;Q9R1Q8 |

**Table S4:**
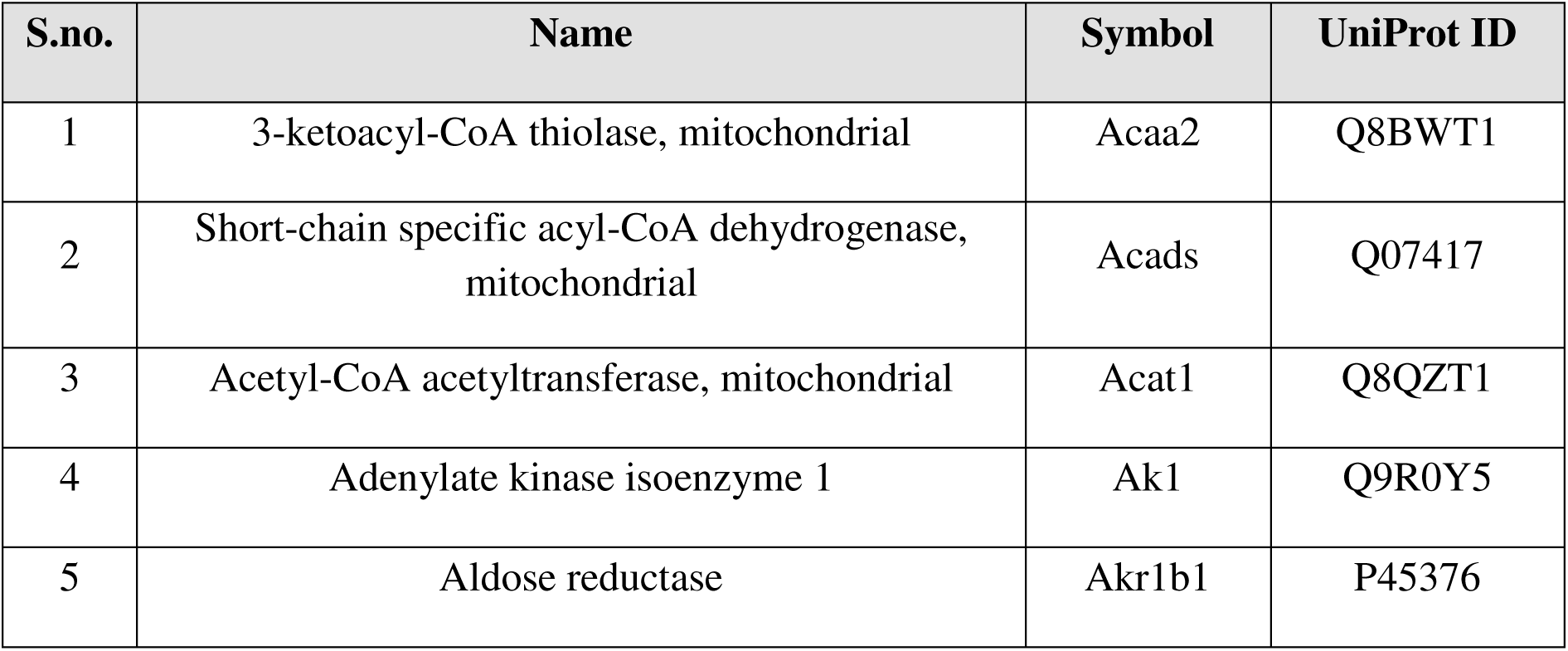
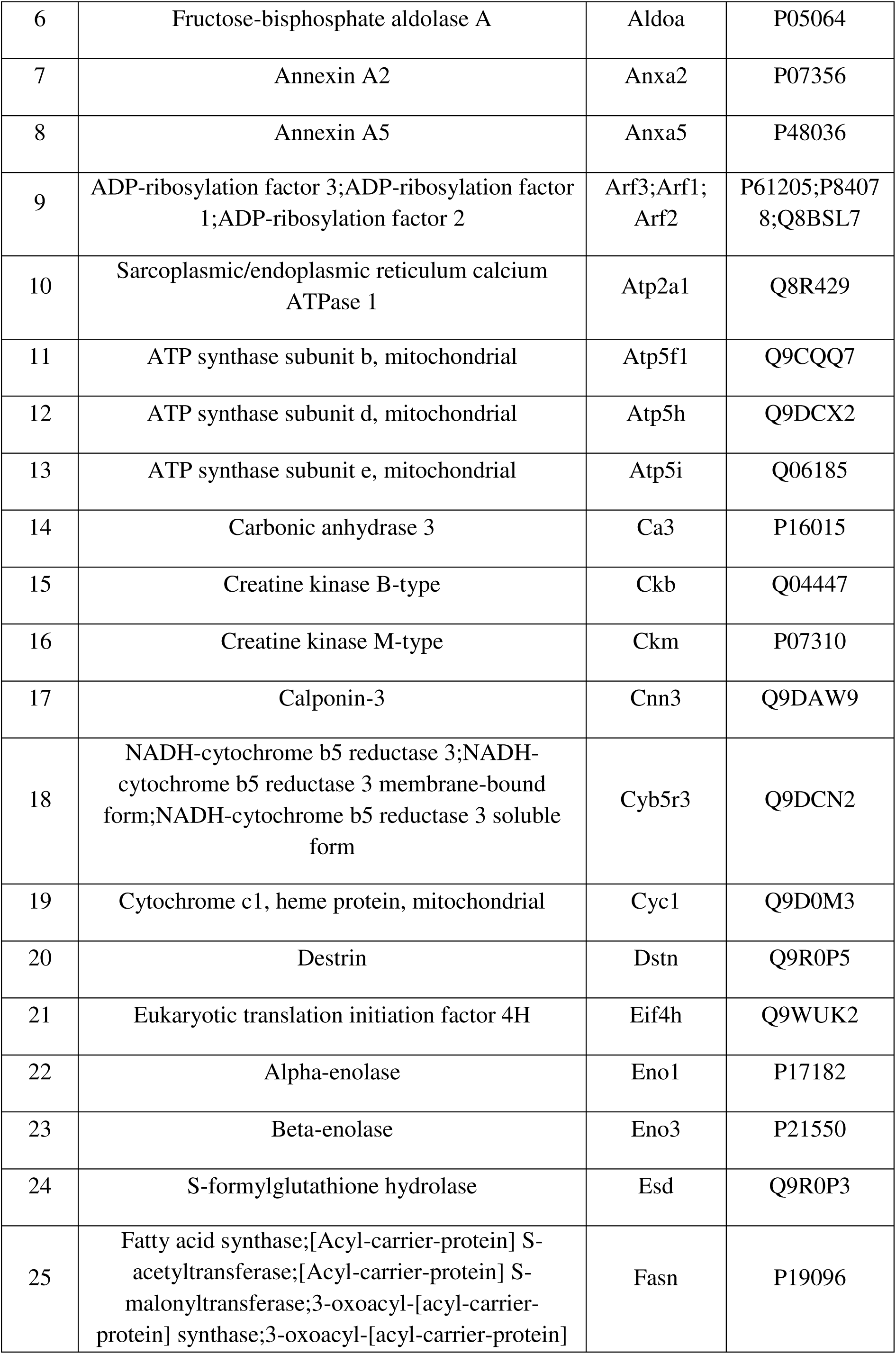
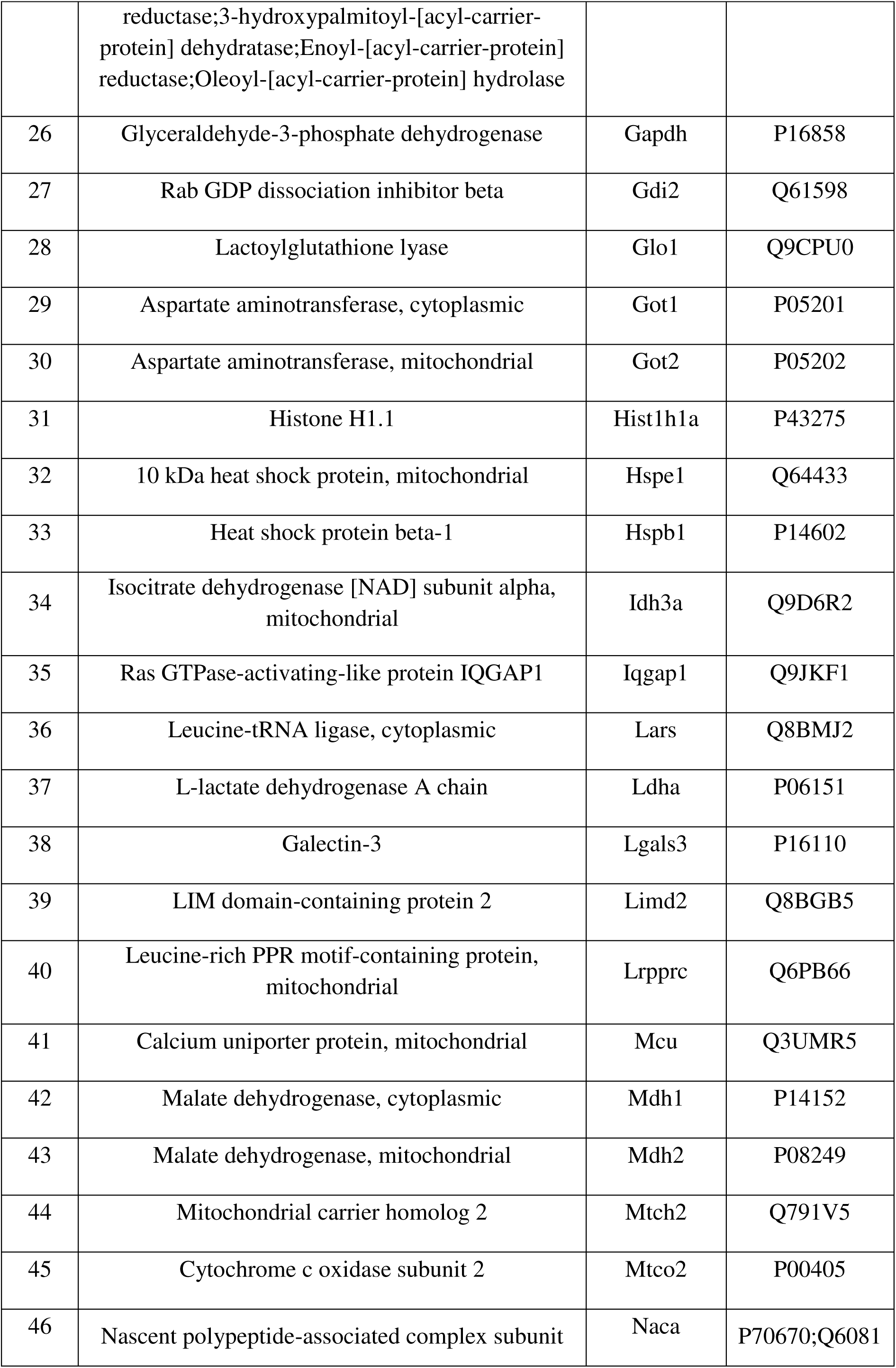
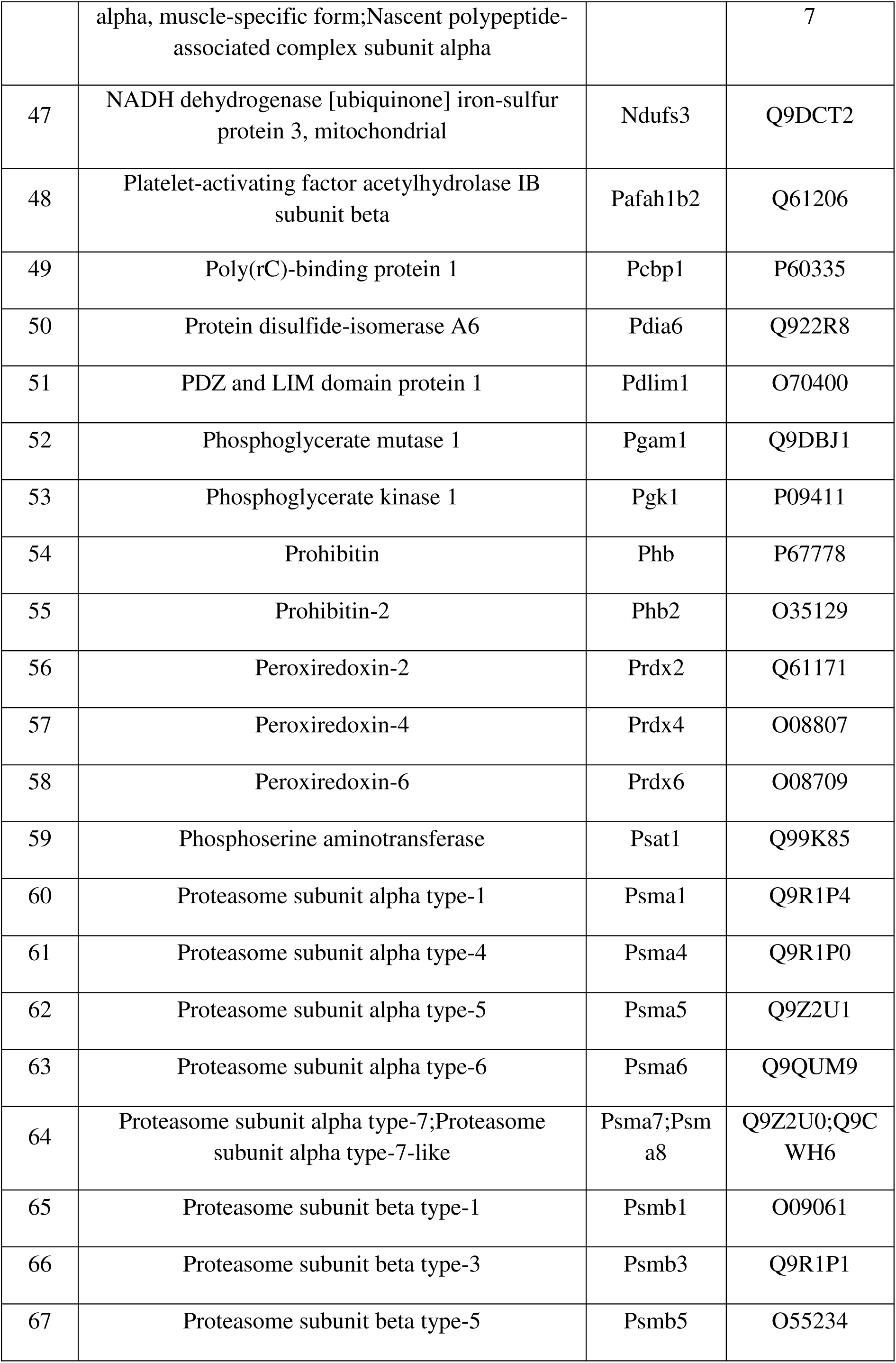
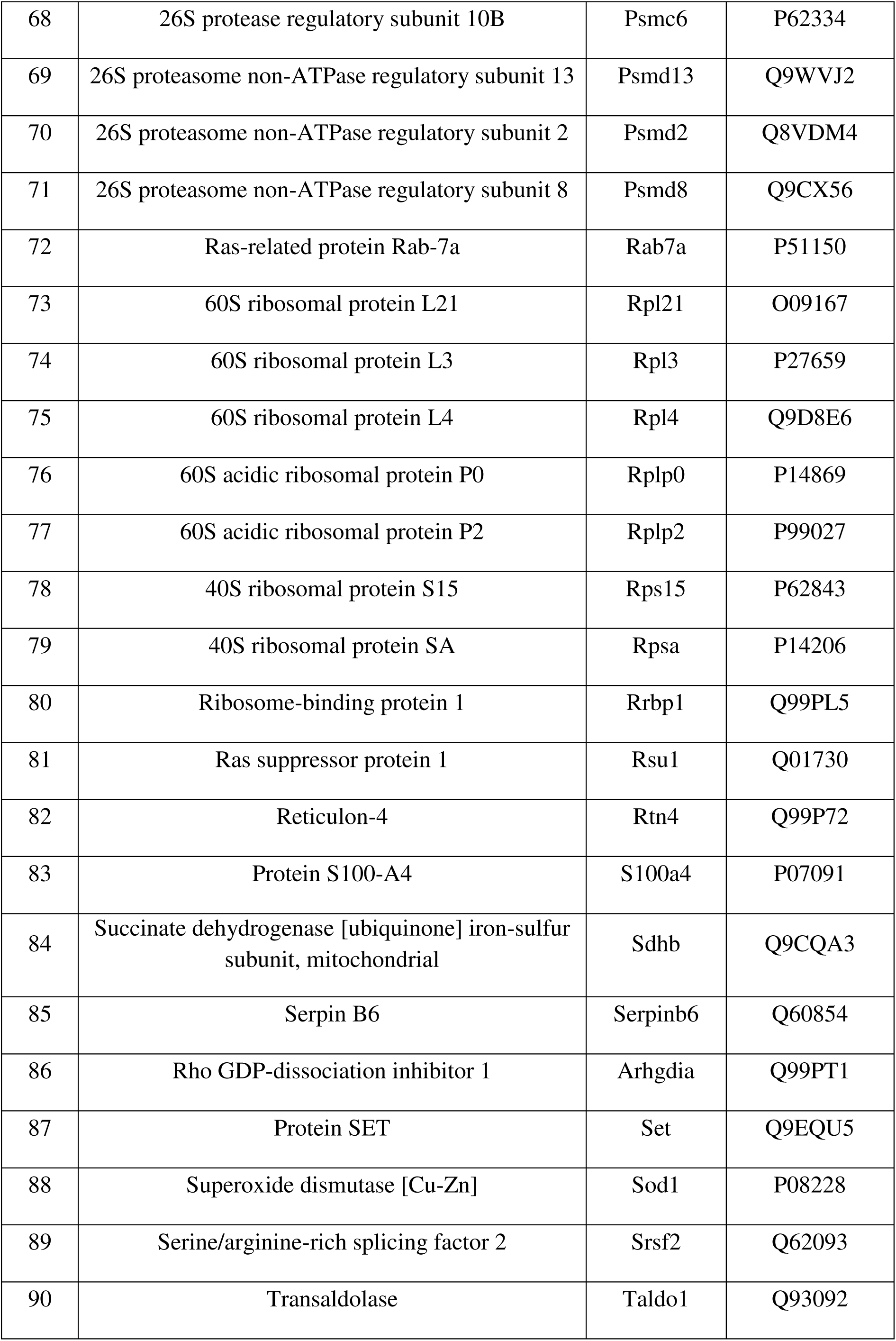
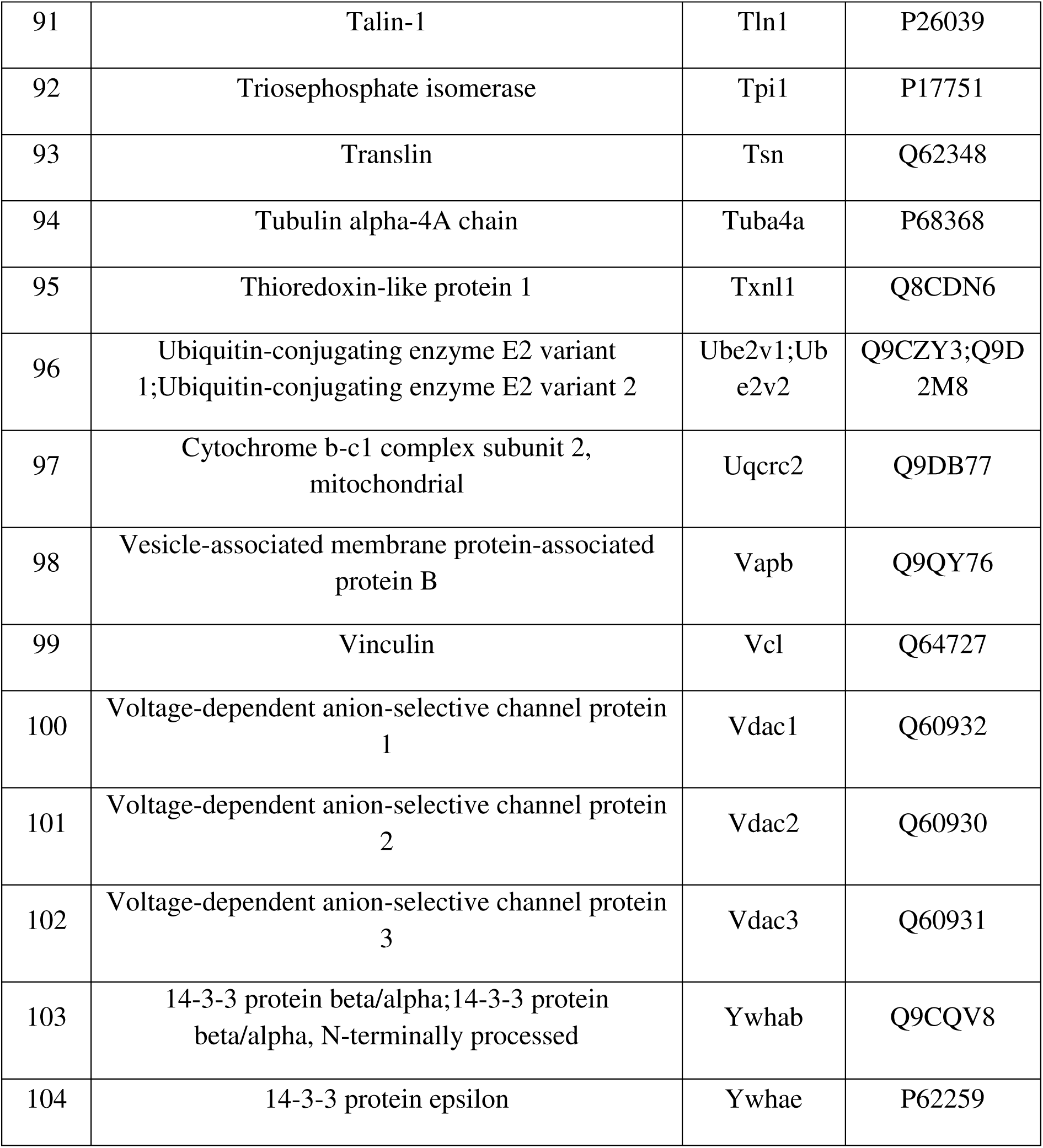
mDia1-interacting proteins identified specifically in MT by LC-MS/MS.

**Table S5:** Interaction Domains on Phb2 and their functional relevance.

| Phb2 Region (aa) | Interacts with | Functional significance | Reference |
| --- | --- | --- | --- |
| 120-232 | Akt2 | Promotes MyoD transactivation function | (Sun et al., 2004) |
| 120-232 | MyoD | Inhibits MyoD transactivation function | (Sun et al., 2004) |
| 175-198 | Estrogen receptor $\alpha$ (ERα) | Represses ERα – mediated transcription | (Delage-Mourroux et al., 2000) |
| 180-232 | mDia1 | Regulates MyoG expression | This study |

**Table S6:** Antibodies used in this study.

| Antibody | IP (μg) | Dilution (IF) | Dilution (Western blot) | Type | Cat. number | Company |
| --- | --- | --- | --- | --- | --- | --- |
| mDia1 | 3 | - | 1:1000 | Monoclonal | 610849 | BD transduction laboratories |
| mDia1 | - | 1:400 | - | Polyclonal | - | Raised in lab against mDia1ΔN3 |
| Phb2 | 3 | - | - | Polyclonal | ab15019 | Abcam |
| Phb2 | - | - | 1:20,000 | Polyclonal | sc-67045 | Santa Cruz |
| Phb2 | - | 1:500 | - | Polyclonal | LS-C287526 | LSBiologicals |
| Phb2 | - | 1:100 | - | Monoclonal | H00011331-M02 | Novus biologicals |
| Akt1 | - | - | 1:1000 | Polyclonal | 2967L | Cell signalling technology |
| Akt2 | - | - | 1:10,000 | Polyclonal | 3063S | Cell signalling technology |
| pAkt2(ser474) | - | - | 1:1000 | Polyclonal | 8599S | Cell signalling technology |
| MyoD | - | 1:100 | 1:1000 | Monoclonal | M3512 | Dako |
| β-actin | - | - | 1:500 | Polyclonal | ab8227 | Abcam |
| GAPDH | - | - | 1:2000 | Monoclonal | ab9484 | Abcam |
| MyoG | - | 1:200 | 1:1000 | Monoclonal | sc-12732 | Santa Cruz |
| Flag | 3 | - | 1:4000 | Monoclonal | F3165 | Sigma |
| Flag | - | 1:500 | - | Polyclonal | F7425 | Sigma |
| GFP | - | - | 1:4000 | Polyclonal | ab6556 | Abcam |
| GFP | - | 1:200 | - | Polyclonal | A10262 | Thermofisher Scientific |
| Active β-Catenin | - | - | 1:1000 | Monoclonal | 05-665 | Millipore |
| Phb1 | - | - | 1:1000 | Polyclonal | 2426S | Cell signalling technology |
| Calreticulin | - | 1:25 | - | Monoclonal | ab22683 | Abcam |
| Cytochrome c | - | 1:500 | - | Monoclonal | 556432 | BD transduction laboratories |
| LaminA/C | - | - | 1:5000 | Polyclonal | ab58529 | Abcam |
| Lamin B1 | - | - | 1:5000 | Polyclonal | ab16048 | Abcam |
| GM130 | - | 1:1000 | - | Monoclonal | 610823 | BD transduction laboratories |
| Peroxidase anti-mouse | - | - | 1:5000 | Polyclonal | 115-035-166 | Jackson ImmunoResearch |
| Peroxidase anti-rabbit | - | - | 1:5000 | Polyclonal | 711-035-152 | Jackson ImmunoResearch |
| Rabbit IgG | 3 | - | - | Polyclonal | 12-370 | Millipore |
| Mouse IgG | 3 | - | - | Polyclonal | 12-371 | Millipore |
| Alexa fluor anti-rabbit 568 | - | 1:500 | - | Polyclonal | A10042 | Thermofisher Scientific) |
| Alexa fluor mouse 647 | - | 1:500 | - | Polyclonal | A-31571 | Thermofisher Scientific |
| Alexa fluor chicken 488 | - | 1:1000 | - | Polyclonal | A11039 | Thermofisher Scientific |

